# Voluntary eating of saltier food by mice and acute stress each abrogate reductions in a neuroinflammatory marker across sexes

**DOI:** 10.1101/2025.07.20.665807

**Authors:** JN Beaver, MT Ford, AE Anello, AK Hite, LM Gilman

**Affiliations:** Department of Psychological Sciences, Kent State University, Kent, OH 44242 USA; Brain Health Research Institute, Kent State University, Kent, OH 44242 USA; Healthy Communities Research Institute, Kent State University, Kent, OH 44242 USA

**Author notes:** Corresponding author: Lee M. Gilman, Ph.D., 600 Hilltop Dr., 144 Kent Hall, Kent State University Kent, OH 44242 USA.

## Abstract

Both consuming excess salt (NaCl) and experiencing environmental stress can elevate neuroinflammation and enhance risk for non-communicable diseases. Most rodent studies investigating these topics use only males, and assess salt or stress separately. Here, we used adult female and male mice to investigate how the combination of access to food high in salt (4% NaCl, w/w) and experiencing an acute stressor interact to affect levels of a proxy measure for neuroinflammation (Iba1). We hypothesized eating salty food and experiencing stress would each individually augment neuroinflammation, and their combination would be additive. Further, we anticipated salty food consumption would increase active stress coping behaviors, and that all of these effects would be enhanced in female mice. Over 4 or 8 weeks, we further evaluated how mice responded to choice access to low (0.4%) and high salt food simultaneously. Our hypothesis that mice across sexes would eventually prefer high over low salt food was supported, while our expectations regarding neuroinflammation and stress did not consistently align with our findings. Instead, we found modest changes in passive coping behaviors driven by our choice condition, unanticipated reductions in sham stress neuroinflammation by high salt in brain region- and biological sex-specific patterns after 4 weeks, and distinct sex- and salt-selective increases in swim stress neuroinflammation after 8 weeks. Though some of our results were unexpected, they include multiple novel and translationally relevant outcomes. Mice willingly choose to eat saltier food over time, akin to people, and this could sex-specifically decrease (females) or augment (males) passive coping stress strategies while eliciting distinctive stress- and brain region-dependent neuroinflammatory patterns over time. Future studies implementing more complex behavior tests and stress manipulations will advance identification of the hidden ways through which salty food and stressful experiences interact to affect risk for non-communicable diseases.

## Introduction

Excess salt (NaCl) intake is an established physiological modulator that can increase risk for several non-communicable diseases (NCDs) including cardiovascular diseases ^1^, obesity ^2,3^, kidney disease ^4^, cancers ^5–7^, type II diabetes ^8,9^, stroke ^10^ and stroke mortality ^11^, dementia ^12,13^, Parkinson’s ^14^, and multiple sclerosis ^15^. A less studied physiological effect of consuming excess salt is neuroinflammation. Microglial activation, indexed by labeling of ionized calcium-binding adaptor molecule 1 (Iba1), can serve as a proxy indicator of salt-elicited neuroinflammation ^16–21^, but no studies have yet analyzed sex as a factor ^17,19,21^. While the ability to study neuroinflammation in humans is quite limited, peripheral inflammatory markers are implicated in the development of/risk for numerous NCDs ^22–29^.

Another contributor to markers of inflammation in people ^30–34^, and neuroinflammation in rodents (though most studies use only males) ^35–38^, is environmental stress. Environmental stress can exacerbate symptoms of, and increase risk for, a range of NCDs ^39–43^. Comorbidity between various NCDs, particularly those affecting the brain and the body (e.g., overweight/obesity and depression symptoms ^44–46)^ may be, in part, facilitated by excess salt intake augmenting the effects of environmental stress and neuroinflammation.

Compared to the expanse of murine papers assessing how high salt intake affects mammalian physiology (for excellent review, see ^47^), relatively few studies have examined how excess salt intake behaviorally alters stress responses, and hardly any have included females (reviewed in ^48^). Murine studies facilitate rigorous control and accurate quantification of salty food consumption, and enable precise measurements of neuroinflammatory and behavioral stress markers in the same individuals. Such invasive manipulations and examinations are at the very least challenging and ethically questionable to perform in humans (explained in depth in reviews ^49–51)^. Thus, while human investigations are unequivocally informative, initial and foundational basic murine research helps focus subsequent scientific questions to be queried studying people.

Our goals here were to advance understanding of how eating food with high or low salt content affects behavioral stress coping responses to an acute stress and subsequent neuroinflammation, including females plus a dietary choice component to expand the translatability of our findings. We evaluated swim stress coping ^52,53^ because different behavioral coping strategies utilized in the face of acute environmental stress have been directly tied to cardiovascular characteristics in male rats ^54^. Additionally, we’ve found that the brain-penetrant anti-inflammatory minocycline mitigates salt intake-induced enhanced active coping plus elevated Iba1 labeling in two stress-responsive brain regions, the paraventricular nucleus of the hypothalamus (PVN) and the basolateral amygdala (BLA) ^16^. Lago and colleagues recently reported that adolescent rats eating salty food (8.0% NaCl) exhibited increased active coping strategies across sexes ^55^. We similarly used female and male mice here. Historically, dietary research utilizing specific chow formulations has restricted rodents to a single salt formulation. To retain precise control over dietary composition while introducing a translational food choice component, we explored diet consumption patterns with chows containing two salt levels (0.4% NaCl, w/w, low salt (LS); or 4.0% NaCl, w/w, high salt (HS)), with or without choice, over shorter (4 week) or longer (8 week) time periods. We investigated how swim stress interacted with chow salt levels to influence behavioral stress coping strategies and Iba1 positive (Iba1^+^) cell numbers in the medial prefrontal cortex, (mPFC; another stress-responsive brain region), PVN, and BLA in the same mice ^56,57^ (see reviews ^58–61)^.

Based on previous evidence ^16–18,20,55,62–64^, we hypothesized male and female mice with access to only HS diet for 4 wks would exhibit enhanced active coping strategies and increased Iba1^+^ counts in the mPFC, PVN, and BLA, relative to male and female mice that only had access to LS diet. Considering reports comparing different salt manipulation durations (8.0% NaC in food, ^65^; 4.0 or 8.0% NaCl in food plus 1.0% NaCl in drinking water, i.e., salt loading, ^21,66^), we anticipated effects detected at 4 wks would be enhanced after 8 wks. Because female rodents can exhibit elevated hormonal and behavioral markers of stress ^67–71^ (but see ^72–74)^, we hypothesized female coping behaviors and Iba1^+^ levels would be higher than males across time points. Lastly, as salt consumption is an innate, genetically-driven mammalian behavior ^75,76^, and salt enhances food palatability ^77–79^, we expected mice with choice access would choose to eat more HS versus LS chow.

## Methods

### Mice

Adult (≥ 9 wks old) male and female C57BL/6J were used for all experiments. Mice were purchased from Jackson Laboratory (Bar Harbor, ME), bred in-house, and group-housed (2-3 mice per cage) in cages containing Nestlets (Ancare, Bellmore, NY), huts (Bio-Serv, Flemington, NJ), and 7090 Teklad Sani-chip bedding (Envigo, East Millstone, NJ). Housing rooms were kept on a 12:12 light:dark cycle, maintained at 22 ± 2°C and 40 ± 10% relative humidity. Experimental mice were fed LabDiet 5001 rodent laboratory chow (LabDiet, Brentwood, MO) *ad libitum* prior to experiment start, and continuing for one week prior to commencing diet manipulations (week 0), to quantify baseline food consumption.

Measurements of food and water consumption, and body weight, occurred twice per week. After week 0, cages were assigned using systematic randomization to diet manipulation durations (i.e., “time period”) of either 4 or 8 wks, and further to one of three diet manipulation conditions: 1) LS only (D17012, Research Diets, Inc., New Brunswick, NJ); 2) HS only (D17013, Research Diets, Inc.); 3) free access to both LS and HS chows (i.e., “mixed”), to analyze voluntary consumption preferences. All mice had *ad libitum* access to their assigned diet(s) and to tap water. All procedures adhered to the National Research Council’s Guide for the Care and Use of Laboratory Animals, 8^th^ Ed. ^80^ and were approved by Kent State University’s Institutional Animal Care and Use Committee.

### Swim Stress

After 4 or 8 wks of diet manipulation, mice either underwent a 6 min swim stress or a sham stress (i.e., no swim stress) condition. Each cage included at least one mouse assigned to each stress condition. Mice were first relocated to individual transport cages, then brought to a holding room adjacent to the testing room 1 h before swim stress. Transport cages contained assigned diet(s) and water, but cage consumption measurements concluded just prior to this short-term individual housing. Procedures of the swim stress are available in the open access publication by Weber and colleagues ^81^. Videos were recorded in black and white to ensure the coder remained unaware of diet condition. Sham stress mice were moved from the holding room directly to the heating pads, and stayed half-on those for 15 min to mimic the experience of swim stress mice, sans swim.

### Stress response behavior scoring

Swim stress behaviors were coded manually offline using the Solomon Coder software program (Solomon Coder v. beta 19.08.02; https://solomon.andraspeter.com). The coder was not informed of diet condition, time period, nor sex, though the latter could be determined through careful visual evaluation. Broadly, these behaviors are generally categorized as active coping (i.e., ambulatory efforts to escape or avoid stressor) and passive coping (i.e., energy conservation to endure or outlast stressor) ^52,53^. Active coping behaviors were defined as swimming and climbing; swimming was characterized as movement of the legs beyond that required to float, and climbing was characterized as taking a vertical posture in the water and activity of all four legs. Passive coping behavior was defined as immobility; immobility was defined as minimum necessary movement of the legs required to float without forward propulsion. Latency to the first immobility bout was measured to indicate the first transition from active to passive coping responses.

### Immunohistochemistry

Two h after undergoing swim stress or sham stress, mice were deeply anesthetized with sodium pentobarbital (FatalPlus; Med-Vet International, Mettawa, IL). After confirming anesthetic plane depth by absences of eye blink and bilateral toe pinch reflexes, mice were perfused transcardially with ice cold 0.9% saline for 3 min followed by ice cold 4.0% paraformaldehyde (PFA) in 0.1 M sodium phosphate buffered saline (PBS) for 7 min, at a rate of 5 mL/min. Brains were extracted and post-fixed in 4% PFA overnight, then transferred to 30% sucrose in 0.1 M PBS for at least 24 h until sufficiently dehydrated. Brains were next coronally sliced at 30 μm on a freezing microtome. Sections were stored in cryoprotectant in 0.1 M sodium phosphate buffer (NaPB) at 4°C until immunohistochemistry (IHC) processing. Cryoprotectant was composed of 20% sucrose (w/v), 30% ethylene glycol (v/v), and 3 mM sodium azide in 0.1 M NaPB.

Immunohistochemistry processing started by rinsing brain sections 3 times for 10 minutes in 0.1M phosphate buffer (PB). Sections then were incubated in blocking solution (0.1M NaPB; 0.9% saline, w/v; and 3% goat serum, v/v (Invitrogen, Waltham, MA)) for 30 min. Next, sections were transferred to 1.5 mL microcentrifuge tubes containing 1:2000 concentrations of the primary antibody in blocking solution (rabbit anti-Iba1; FUJIFILM Labchem Wako 019-19741, Osaka, Japan) and left on a horizontal shaker at room temperature for 48 h. Following primary incubation, brain sections were again rinsed 3 times in 0.1M NaPB before undergoing a 2 h incubation in 1:250 biotinylated goat anti-rabbit IgG secondary antibody (Invitrogen A16108, Waltham, MA) in blocking solution. After 3 more 0.1M NaPB rinses, sections were incubated for 10 minutes in 1:250 streptavidin-AlexaFluor 488 conjugate (Invitrogen S11223) in 0.1M NaPB. Finally, sections were rinsed three times in 0.1M NaPB, then briefly submerged in deionized water for mounting on 25mm × 75mm × 1.0mm Fisherbrand™ Superfrost™ Plus Microscope Slides (Fisher Scientific 12-550-15, Hampton, NH) using 24mm × 60mm GOLD SEAL Cover Glass (Electron Microscopy Sciences 63770-0, Hatfield, PA) and ProLong Diamond Antifade Mountant (Invitrogen P36971), and left to dry overnight. The next day, clear nail polish was applied to seal all coverslip edges.

Images of immunolabeled brain sections were obtained using an Olympus FluoView 3000 confocal microscope with 20x magnification. The software program ImageJ (NIH, Bethesda, MD) was used to quantify Iba1^+^ cells ^82^. Each image was despeckled to filter noise. Next, background was subtracted using a 30 pixel threshold. The triangle auto threshold option was selected ^83–86^, then images were converted to a binary image. Counts of Iba1^+^ within the mPFC, PVN, and BLA were made in each hemisphere when possible, and no significant differences were found between hemispheres. Parameters for infinity (15) and for circularity (0.5-1) were kept consistent across images. The image coder was not informed of diet condition, time period, stress condition, nor sex.

### Statistical Analyses

Estimated marginal means were calculated using IBM SPSS Statistics 29.0.1.1 (244) (IBM, Armonk, NY) then graphed with GraphPad Prism 10.5.0 (673) (GraphPad Software, San Diego, CA). Our significance threshold was set *a priori* at p<0.05. Estimated marginal means (EM means) ± 95% confidence intervals (CIs) were graphed. Swim behaviors were analyzed across time periods using three-way univariate general linear models (GLMs; diet × sex × time period). Iba1^+^ cell counts were analyzed within each brain region and time period using three-way univariate GLMs (diet × sex × stress condition). Adult mice naturally exhibit sexual dimorphism in body weights, with males weighing more than females ^87,88^. Therefore, consumption per day of food, salt, and water was normalized to cage body weight to facilitate cross-sex comparisons. Week 0 normalized consumption of food and water were analyzed with three-way univariate GLMs (future diet × sex × time period) to confirm no detectable confounds existed prior to diet manipulation. Individual mouse body weights, and normalized cage food, salt, and water per day consumption rates were each analyzed within time period using 2-way repeated measures GLMs (repeated measure × diet × sex). Greenhouse-Geisser corrections were used for within-subjects GLM analyses. All graphed data were also analyzed with pairwise comparisons using Bonferroni correction. Non-significant trends with p<0.10 but p>0.05 were only discussed if the corresponding partial η^2^ (η_p_^2^) ≥0.06 ^89^.

## Results

Body weights of individual mice were not affected by any three-nor two-way interactions with diet (Table S1). A significant two-way interaction of week × sex for body weight was detected for mice in the 8 wk time period, whereas for the 4 wk time period only main effects of time and sex were significant (Table S1). Pairwise comparisons revealed no differences between diet at any week for female mice across time periods (Fig. S1A, C). At most weeks, male mice across time periods also exhibited no differences in body weight as a result of diet (Fig. S1B, D). Only at weeks 1 and 2 did 4 wk mixed male mice weigh less than LS males (Fig. S1B); and only at week 8 did mixed 8 wk males weigh more than HS males (Fig. S1D). As expected, pairwise comparisons indicated that males in both time periods and on all diets weighed more than females in the same time period and diet at every single week (Fig. S1).

Baseline, or week 0, normalized daily food and water consumption did not exhibit any significant three-nor two-way interactions involving time period or diet (Table S3). This illustrates that baseline normalized food and water consumption rates likely did not confound measures of same during diet manipulation. For both normalized food/day and water/day, the only significant main effect was sex (Table S3). Normalized salt/day could not be analyzed for week 0 because the NaCl content of LabDiet 5001 is not disclosed and is considered proprietary.

Following week 0, cages of same sex mice were assigned to one of three diets conditions for either 4 or 8 wks before undergoing swim or sham stress, then perfusions 2 h later for subsequent Iba1^+^ labeling. Normalized daily food consumption was analyzed on a per week basis. No significant three-way interactions were found for week × sex × diet for either time period (Table 1). For both the 4 and 8 wk time periods, significant interactions between week × diet were observed (Table 1). A significant main effect of sex was detected for the 8 wk, but not the 4 wk, time period (Table 1). Considering these were analyses of normalized food consumption to account for body weight sexual dimorphism, the main effect of sex in the 8 wk time period was not anticipated. Considering this main effect of sex in conjunction with the visualized data (Fig. 1), and that it only occurred in the 8 wk time condition, it is likely that the extended time period facilitated detection. Further, given these were normalized data, the sex effect must be attributable to something other than body weight. Pairwise comparisons found 4 wk HS females ate more at weeks 3 and 4 than both LS females (Fig. 1A) and HS males (Fig. 1A, B). These findings were not duplicated in the 8 wk time period (Fig. 1C, D). Rather, 8 wk HS males ate more than 8 wk LS males only at week 6 (Fig. 1D), and 8 wk mixed females ate more than 8 wk mixed males only at week 4 (Fig. 1C, D).

**Figure 1.**
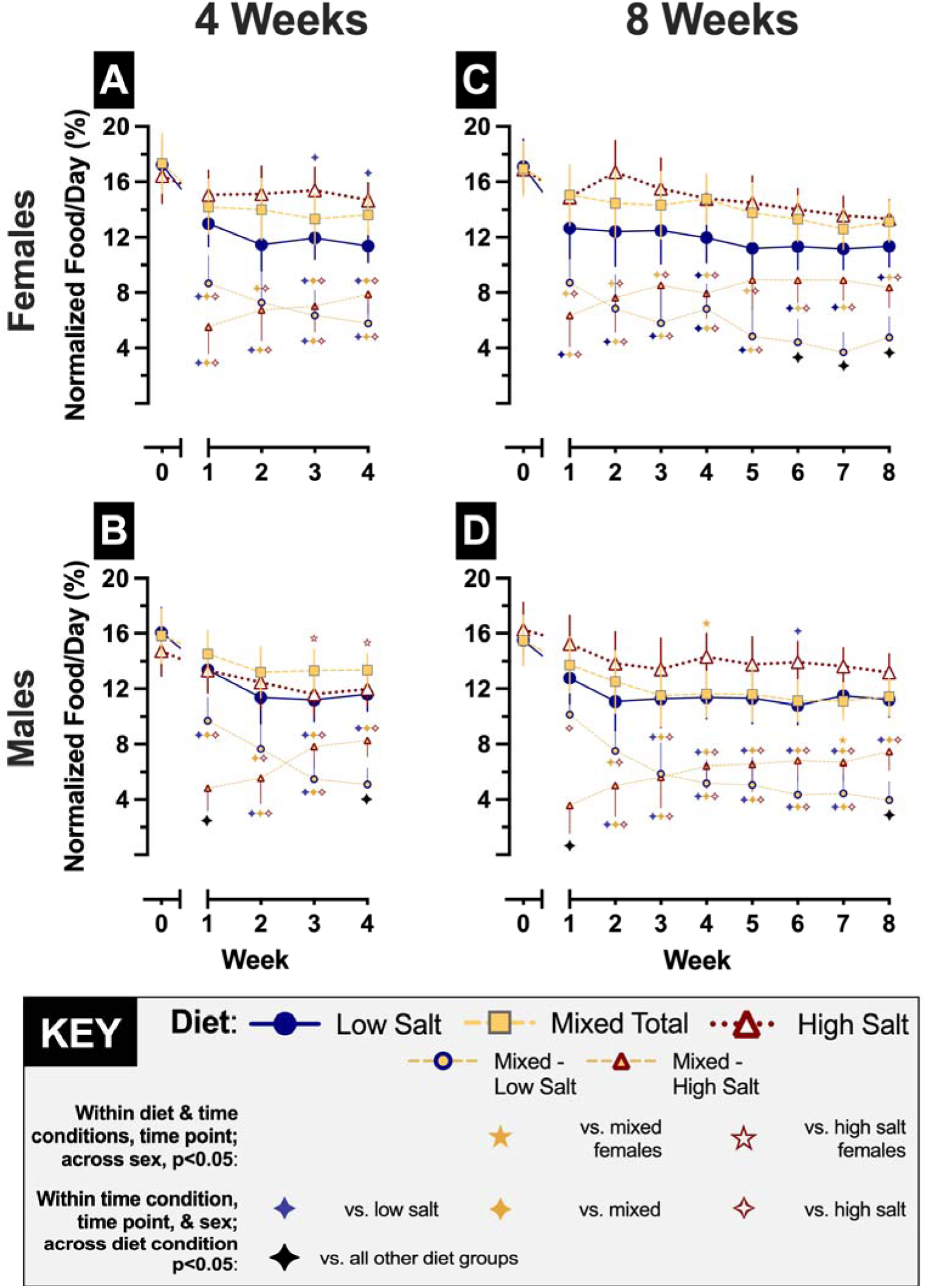
Weekly rates of food consumed per day normalized to cage body weight. Normalized food/day intake was calculated weekly for 4 wk (panels A-B) and 8 wk (panels C-D) mice. Female cages are shown in panels A,C; male cages are in panels B, D. Cages assigned to low salt diet (0.4% NaCl, w/w) are shown in blue circles with solid lines. Cages assigned to high salt diet (4.0% NaCl) are displayed in white-filled red triangles with dotted lines. Mixed cages given free choice access to both low and high salt diets are graphed in grey-bordered yellow squares with thick dashed lines for total food consumption. The component of total mixed cage consumption comprised of low salt diet is graphed in yellow-filled blue circles; the component of total mixed cage consumption comprised of high salt diet is graphed in yellow-filled red triangles; both have thin dashed lines. Cage Ns=5-7 within each sex/diet/time period; for exact numbers, see Supplemental Table S4. Data are graphed as EM means ± 95% CI. Blue, yellow, and red diamonds indicate p<0.05 (see Tables S5 and S6 for 4 wk and 8 wk, respectively, exact p values) compared with ⍰low salt, ⍰mixed total, and ⍰high salt cage consumption in same sex at same week within same time period. Black diamonds (⍰) indicate p<0.05 versus all other points within same sex, week, and time period. Left to right, 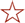 indicates p=0.002, 0.003 vs. high salt females in same week and time period. Left to right, 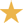 indicates p=0.013, 0.029 vs. mixed total and mixed high salt females, respectively, in same week and time period.

**Table 1.**
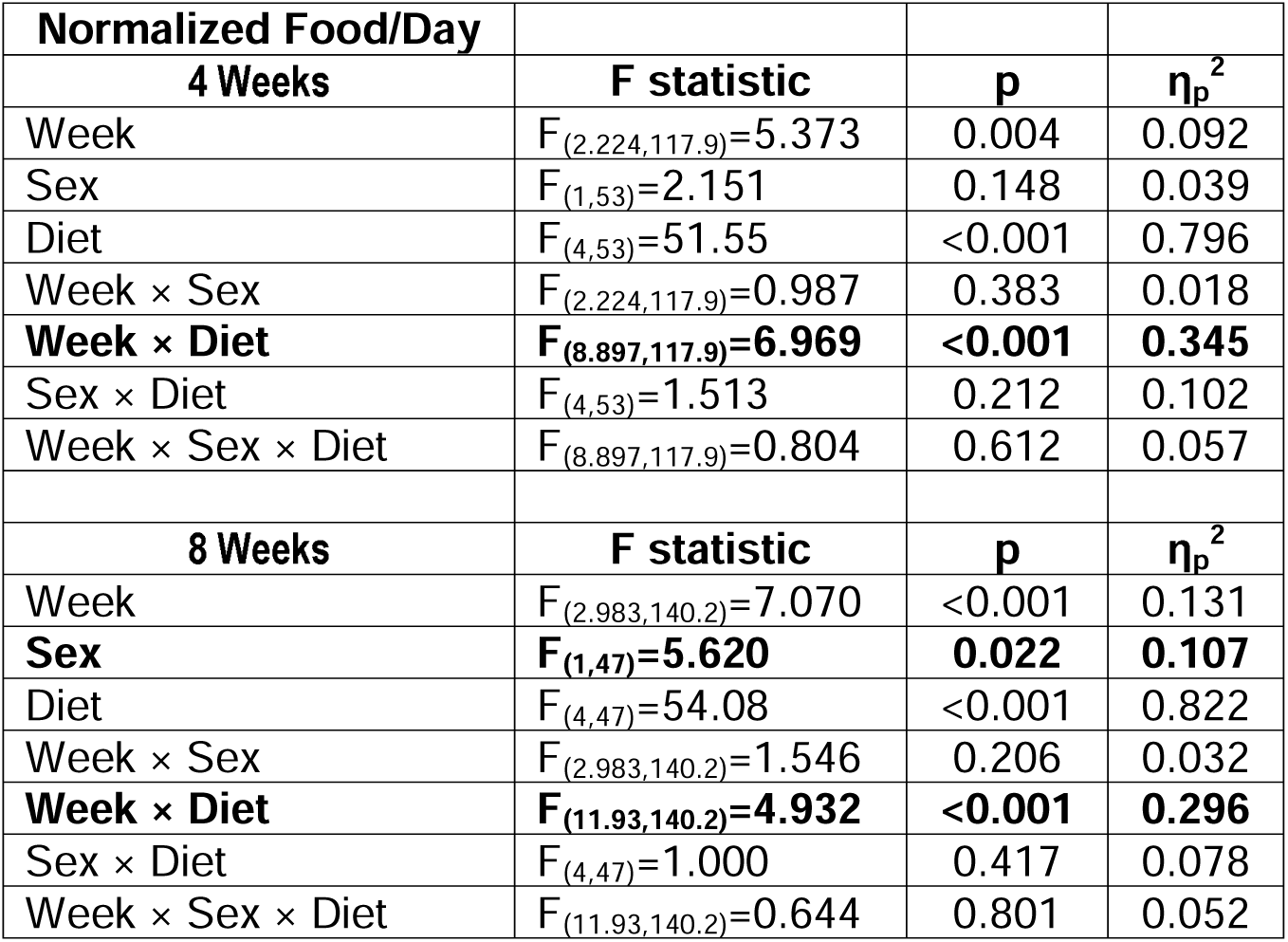
Two-way RM GLMs of normalized food/day.

What was consistently observed across 4 and 8 wk time periods relates specifically to mixed mice. This is the first time that rodents’ choice between two purified diet formulations with LS and HS levels has been evaluated. While single formulation diet studies are essential for empirically determining the contributions of specific food components to physiology and behavior, these are inherently limited in their translatability to people. Continual choice access offered to rodents is a substantial step towards enhancing the translational contribution of our studies. In 4 wk mice, mixed females ate similar levels of LS and HS diets (Fig. 1A). At the start of the 4 wk time period, mixed male mice started by exhibiting a clear preference for LS food, but this reversed by the end (Fig. 1B). Similar, though not identical, diet choices were found in the 8 wk time period. Mixed females once again consumed comparable amounts of LS and HS diets for the first 5 weeks (Fig. 1C). It is not until weeks 6-8 that 8 wk mixed females significantly prefer HS compared to LS diet (Fig. 1C). Further mirroring the 4 wk time period (Fig. 1B), 8 wk mixed males began by consuming significantly more LS than HS chow (Fig. 1D), and this preference disappeared by week 2. In contrast to 4 wk mixed males (Fig. 1B), 8 wk mixed males did not attain a significant preference for HS over LS diet until week 8 (Fig. 1D). At only one week across both time conditions did a sex difference in mixed diet preferences emerge; at week 7, mixed males ate significantly less HS diet than mixed females at the same week (Fig. 1C, D). Critically, total normalized food consumption by mixed mice never differed from same sex LS and HS mice in the same time periods (Fig. 1), illustrating that these choice access consumption patterns are not confounded by significantly more or less overall food intake. Moreover, our choice experiments emphasize how, like humans, mice will voluntarily eat more salty food even when less salty food is available, and despite HS food containing far beyond what mice physiologically require for salt. In fact, mice and rats require as little as 0.05% NaCl, w/w ^90^ [(see Papers for proper full reference)].

Mice adjust their water intake in accord with their salt intake, thereby avoiding hyperosmotic stress even when their only food access is to HS chow ^63^. We again observed such behavior here (Table S7, Fig. S2). Neither the 4 nor 8 wk time periods exhibited significant week × sex × diet interactions (Table S7). Two-way interaction and main effect analyses differed across time periods. In the 4 wk time period, a significant interaction of sex × diet plus a main effect of week was found (Table S7). For the 8 wk time period, it was the week × sex interaction that was significant, as was a main effect of diet (Table S7). Despite these variations, pairwise comparisons illustrated a broadly uniform theme, in that that HS mice of both sexes and across 4 and 8 wk time periods almost always consumed more normalized water than LS mice of the same sex at each corresponding week (sole exception: 8 wk male HS mice at week 2; Fig. S2D). Normalized water consumption in mixed mice was far more variable. This is to be expected both because mice have access to two diets instead of one, and because each cage contained two to three mice. Mixed 4 wk females’ normalized water intake was consistently less than that of HS 4 wk females, and not different from LS 4 wk females (Fig. S2A). The 8 wk females partially replicated this, with mixed females’ normalized water intake being less than HS females’ at weeks 1, 2, and 4 (Fig. S2C). Only at week 1 did mixed 8 wk females drink more water than LS 8 wk females (Fig. S2C). Normalized drinking in mixed males was more discrepant across time periods. Mixed 4 wk males exhibited more normalized water intake than LS 4 wk males at weeks 2-4 (Fig. S2B), whereas mixed 8 wk males never displayed differences in normalized drinking versus LS 8 wk males (Fig. S2D). Such inconsistencies may reflect the related delay in dietary preference switch from LS to HS in 8 wk compared to 4 wk mixed males (Fig. 1B, D). Studies assessing salty food choice and water consumption in singly, rather than group, housed males could directly evaluate this supposition. Regarding sex differences, normalized water consumption by HS 4 wk males was less than HS 4 wk females across weeks 1-3 (Fig. S2A, B). The same direction of this HS sex difference was also found in the 8 wk time period at weeks 1-2 (Fig. S2C, D). Mixed 8 wk males drank less normalized water than mixed 8 wk females only at week 1.

To drill down a bit further into normalized water intake variations, and to pare down to the fundamental salt impact of mixed diet choices compared to LS or HS only diets, we calculated normalized daily salt intake per cage on a weekly basis (Table 2, Figure 2). No significant week × sex × diet interaction was found for either time period (Table 2). Two two-way interactions, of week × diet and sex × diet, were significant for normalized salt intake in 4 wk mice (Table 2).

**Figure 2.**
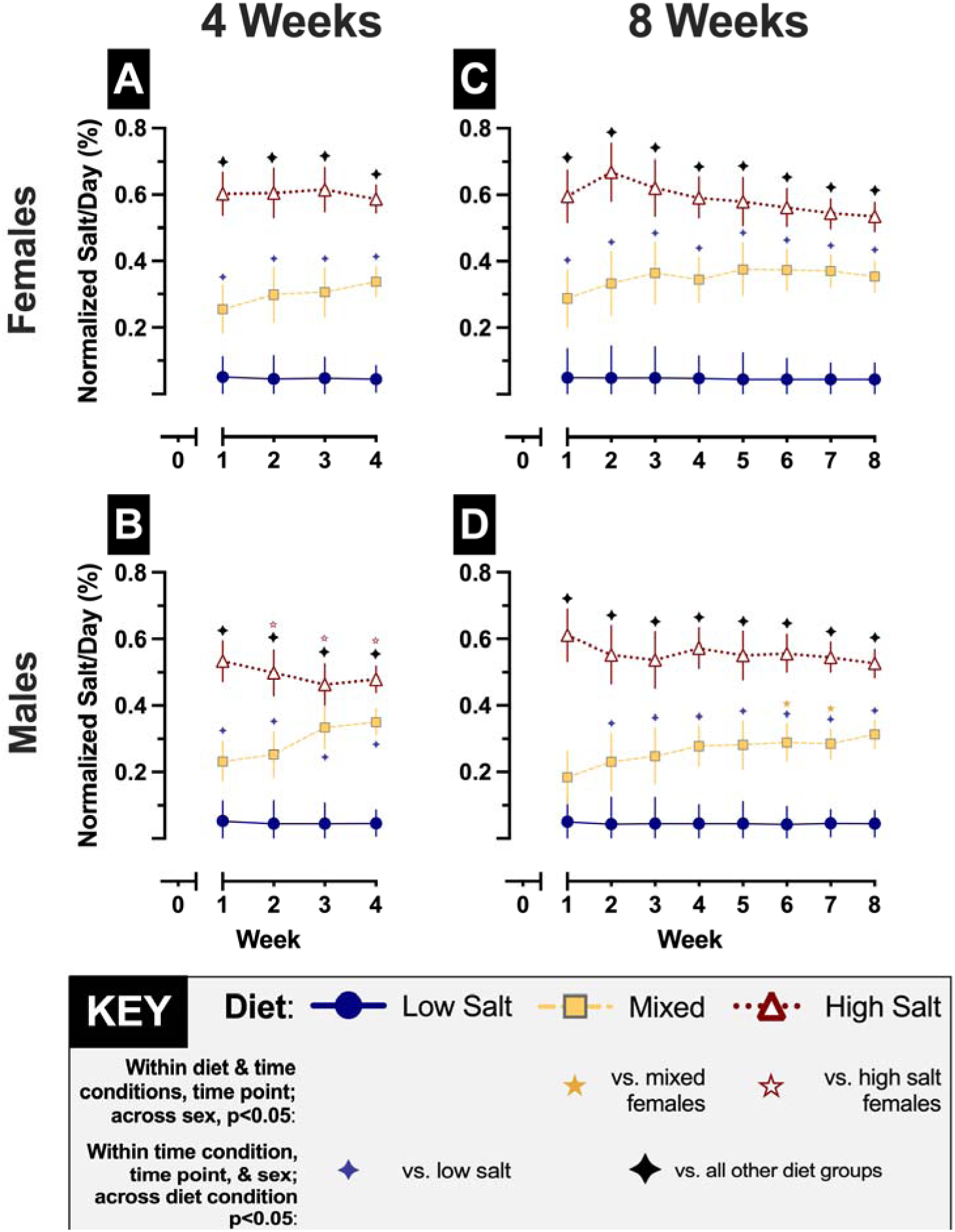
Weekly rates of salt consumed per day normalized to cage body weight. Normalized salt/day intake was calculated weekly for 4 wk (panels A-B) and 8 wk (panels C-D) mice. Female cages are shown in panels A,C; male cages are in panels B, D. Cages assigned to low salt diet (0.4% NaCl, w/w) are shown in blue circles with solid lines. Cages assigned to high salt diet (4.0% NaCl) are displayed in white-filled red triangles with dotted lines. Mixed cages given free choice access to both low and high salt diets are graphed in grey-bordered yellow squares with dashed lines. Cage Ns=5-7 within each sex/diet/time period; for exact numbers, see Supplemental Table S4. Data are graphed as EM means ± 95% CI. For 4 wk time period, blue and black diamonds indicate p<0.001 (except males week 3, mixed vs. high salt, p=0.016) compared with ⍰low salt ⍰and all other diets in same sex at same week within same time period. For 8 wk time period, blue and black diamonds indicate p<0.05 (see Table S10 for exact p values) compared with ⍰low salt and ⍰all other diets in same sex at same week within same time period. Left to right, 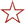 indicates p=0.042, 0.002, <0.001 vs. high salt females in same week and time period. Left to right, 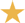 indicates p=0.049, 0.012 vs. mixed females in same week and time period.

**Table 2.**
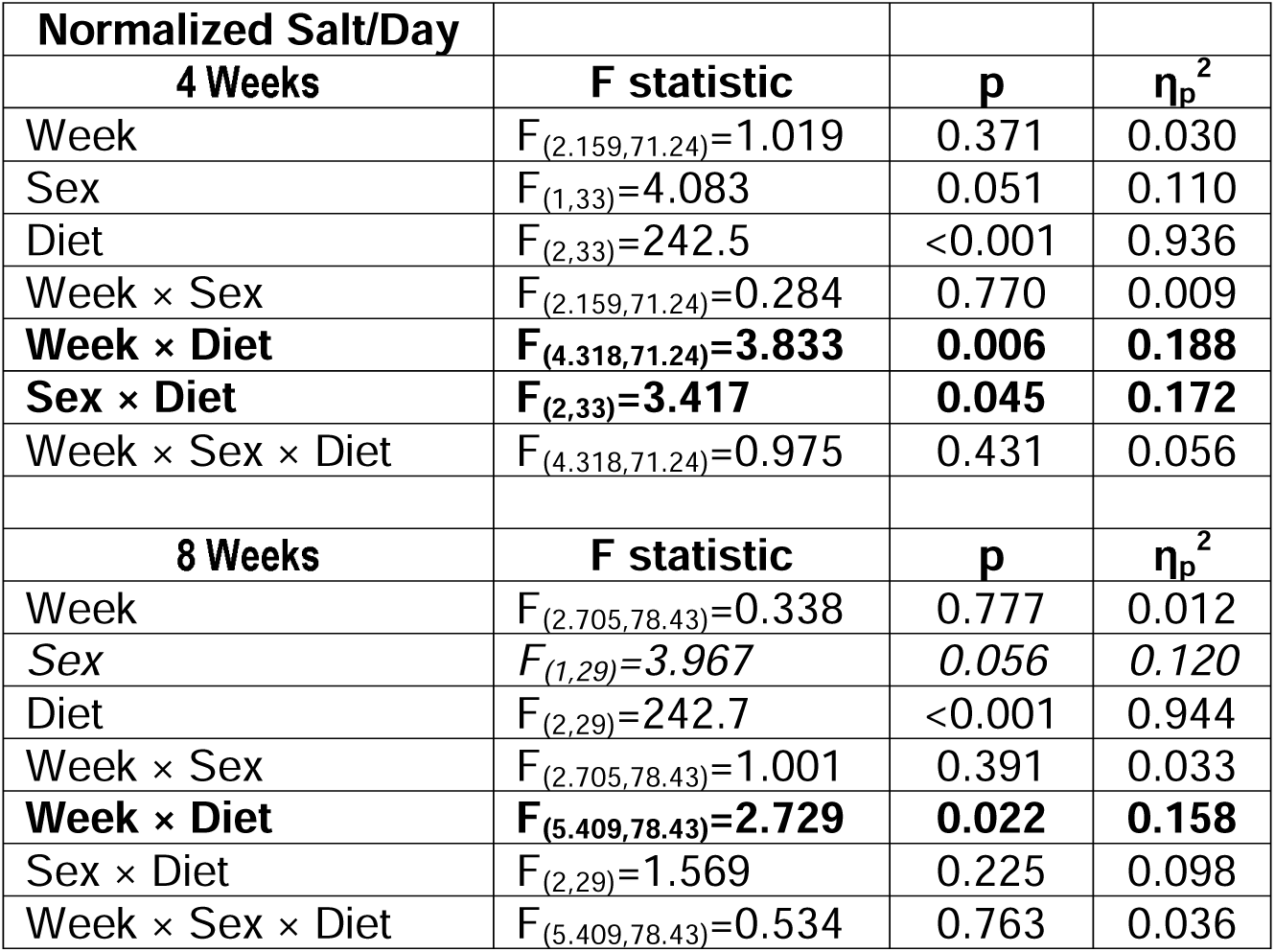
Two-way RM GLMs of normalized salt/day.

While 8 wk mice also had a significant week × diet interaction, this time period exhibited a non-significant trend regarding sex (p=0.056, η_p_^2^ =0.120; Table 2). Pairwise comparisons showed, as expected, across sexes and both time periods, HS mice always exhibited elevated normalized salt intake compared to same sex LS mice at the same week (Fig. 2). With only a single exception (8 wk male mixed mice at week 2; Fig. 2D), mixed mice of all sexes and time conditions also ate more salt than same sex LS mice in the same week (Fig. 2). Largely corresponding to normalized food intake, sex differences in normalized salt intake were found at weeks 2-4 in 4 wk mice, where HS males ate less normalized salt than HS females; and at weeks 6-7 in 8 wk mice, with mixed males consuming less normalized salt than mixed females (Fig. 2).

Immediately after the conclusion of measuring body weights and consumption for 4 or 8 wk diet manipulations, at least one mouse from each cage was assigned to a swim stress, and at least one mouse from each cage was assigned to a sham stress. In contrast to our hypothesis, we did not observe any significant three-nor two-way interactions involving diet on active coping behaviors (swimming and climbing; Table 3, Figure 3A, B). For climbing, we did find a significant sex × time interaction (Table 3). This looked to be driven by climbing increasing in 8 wk males compared to 4 wk males plus 4 and 8 wk females (Fig. 3B). Pairwise comparisons found that both LS and HS males in the 8 wk time period exhibited significantly more climbing than 4 wk males on the same respective diets (Fig. 3B). Passive coping, indexed by immobility, did not exhibit a significant sex × time × diet interaction, but a sex × diet interaction was significant (Table 3). Visually it appears this could be from mixed females in both diet conditions exhibiting less immobility compared to LS and HS females, whereas mixed males in both diet conditions seemed to have more immobility versus LS and HS males (Fig. 3C). Pairwise comparisons did not reveal any significant differences for immobility (Fig. 3C). As with the other swim stress behaviors, latency to the first immobility bout also did not exhibit a significant three-way interaction (Table 3). Significant sex × time and diet × time interactions were observed (Table 3). Only one significant pairwise comparison was observed, with LS 8 wk males displaying a longer latency to their first immobility bout compared to LS 4 wk males (Fig. 3D).

**Figure 3.**
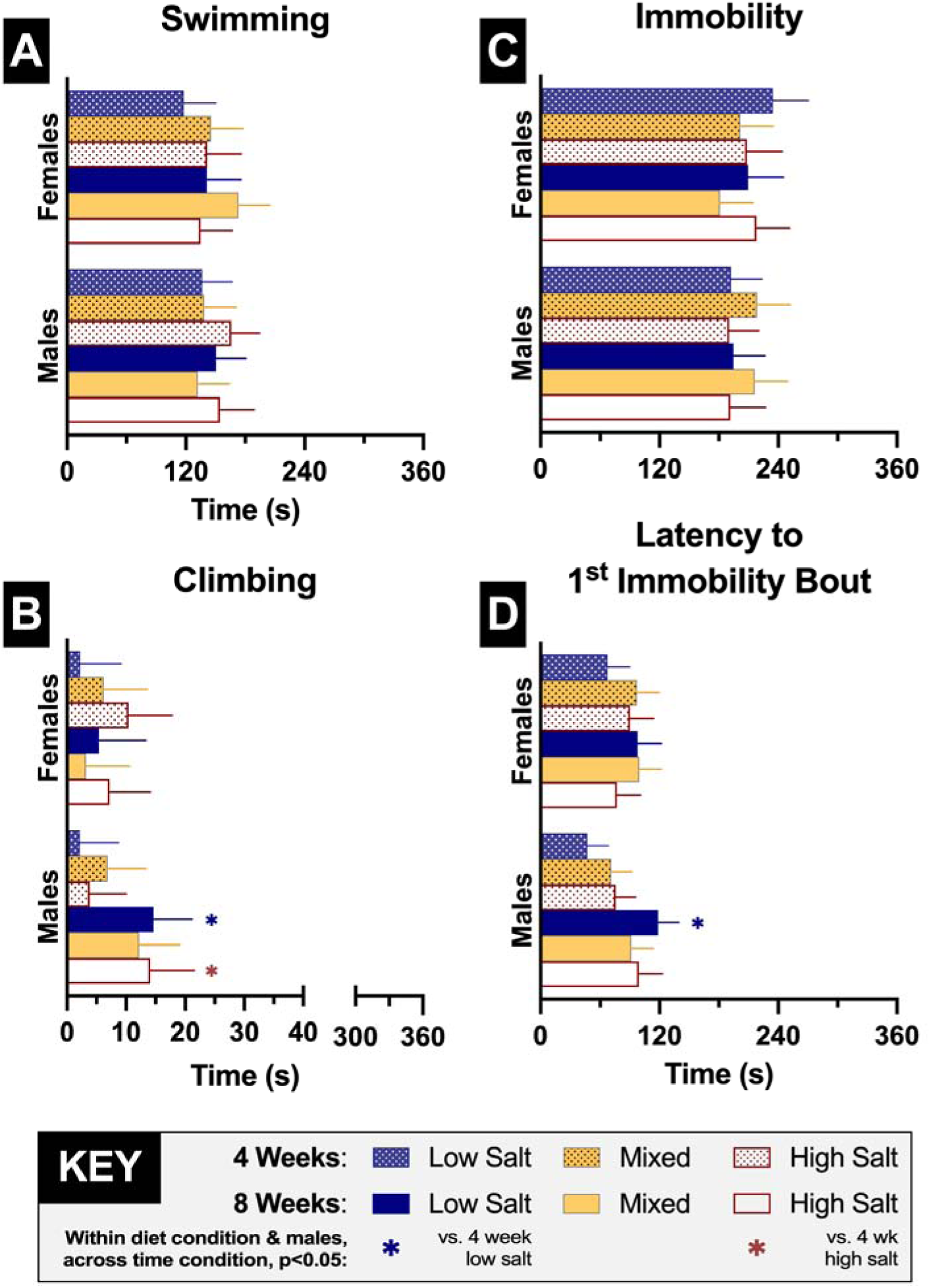
Active and passive coping behaviors during swim stress. Active coping behaviors are graphed in panels A (swimming) and B (climbing). Passive coping behaviors are graphed in panel C (immobility). Initial coping strategy switch from active to passive is in panel D (latency to first immobility bout). Mice assigned to low salt diet (0.4% NaCl, w/w) are shown in blue bars. Mice assigned to high salt diet (4.0% NaCl) are displayed in white-filled red bars. Mixed cages given free choice access to both low and high salt diets are graphed in grey-bordered yellow bars. Dotted bars are for 4 wk data, clear bars are for 8 wk data. Ns=7-10 within each sex/diet/time period; for exact numbers, see Supplemental Table S11. Data are graphed as EM means ± 95% CI. Left to right, ⍰indicates p=0.009, <0.001 vs. 4 wk low salt males. ⍰indicates p=0.037 vs. 4 wk high salt males.

**Table 3.** Two-way RM GLMs of swim stress behaviors.

| **Swim Stress Behaviors** |  |  |  |
| --- | --- | --- | --- |
| **Swimming** | **F statistic** | **p** | **$\eta_p^2$** |
| Sex | $F_{(1,85)}=0.178$ | 0.674 | 0.002 |
| Diet | $F_{(2,85)}=0.716$ | 0.492 | 0.017 |
| Time | $F_{(1,85)}=0.515$ | 0.475 | 0.006 |
| Sex × Diet | $F_{(2,85)}=2.279$ | 0.109 | 0.051 |
| Sex × Time | $F_{(1,85)}=0.722$ | 0.398 | 0.008 |
| Diet × Time | $F_{(2,85)}=0.759$ | 0.471 | 0.018 |
| Sex × Diet × Time | $F_{(2,85)}=0.239$ | 0.788 | 0.006 |
| **Climbing** | **F statistic** | **p** | **$\eta_p^2$** |
| Sex | $F_{(1,83)}=2.449$ | 0.121 | 0.029 |
| Diet | $F_{(2,83)}=0.636$ | 0.532 | 0.015 |
| Time | $F_{(1,83)}=4.168$ | 0.044 | 0.048 |
| Sex × Diet | $F_{(2,83)}=0.555$ | 0.576 | 0.013 |
| **Sex × Time** | **$F_{(1,83)}=6.451$** | **0.013** | **0.072** |
| Diet × Time | $F_{(2,83)}=0.908$ | 0.407 | 0.021 |
| Sex × Diet × Time | $F_{(2,83)}=0.151$ | 0.860 | 0.004 |
| **Immobility** | **F statistic** | **p** | **$\eta_p^2$** |
| Sex | $F_{(1,84)}=0.703$ | 0.404 | 0.008 |
| Diet | $F_{(2,84)}=0.118$ | 0.888 | 0.003 |
| Time | $F_{(1,84)}=0.346$ | 0.558 | 0.004 |
| **Sex × Diet** | **$F_{(2,84)}=3.129$** | **0.049** | **0.069** |
| Sex × Time | $F_{(1,84)}=0.398$ | 0.530 | 0.005 |
| Diet × Time | $F_{(2,84)}=0.329$ | 0.721 | 0.008 |
| Sex × Diet × Time | $F_{(2,84)}=0.311$ | 0.733 | 0.007 |
| **Latency to 1<sup>st</sup> Immobility Bout** | **F statistic** | **p** | **$\eta_p^2$** |
| Sex | $F_{(1,85)}=0.439$ | 0.509 | 0.005 |
| Diet | $F_{(2,85)}=0.397$ | 0.674 | 0.009 |
| Time | $F_{(1,85)}=11.79$ | <0.001 | 0.122 |
| Sex × Diet | $F_{(2,85)}=0.993$ | 0.375 | 0.023 |
| **Sex × Time** | **$F_{(1,85)}=5.910$** | **0.017** | **0.065** |
| **Diet × Time** | **$F_{(2,85)}=4.849$** | **0.010** | **0.102** |
| Sex × Diet × Time | $F_{(2,85)}=0.293$ | 0.747 | 0.007 |

Two h after swim or sham stress, mice were deeply anesthetized then transcardially perfused to facilitate immunolabeling of Iba1^+^ in three stress-responsive brain regions: mPFC, PVN, and BLA. Iba1^+^ cell counts in the brain serve as a proxy index of neuroinflammation ^82^ (but see ^91,92^). Counts of Iba1^+^ cells for 4 wk mice did not exhibit any significant sex × stress × diet interactions in any of the three brain regions (Table 4). No significant two-way interactions were detected in the mPFC, but a main effect of diet was observed (Table 4). Pairwise comparisons reported that, again counter to our hypothesis, HS reduced Iba1^+^ cells compared to LS selectively in sham stress 4 wk females (Fig. 4A). For the PVN, no two-way nor main effects were significant (Table 4). Two non-significant trends for sex × diet (p=0.057, η_p_^2^=0.093) and diet (p=0.083, η_p_^2^=0.081) were noted (Table 4). Pairwise comparisons for 4 wk PVN Iba1^+^ cells demonstrated that sham stress HS males displayed less than LS males in the same stress condition (Fig. 4B). Additionally, both sham stress 4 wk LS females and swim stress 4 wk LS males had lower Iba1^+^ cells than sham stress 4 wk LS males (Fig. 4B). For 4 wk BLA Iba1^+^ cells, no significant interactions nor main effects were detected for stress, sex, and diet (Table 4). Aligning with omnibus statistical analyses, pairwise comparisons revealed no significant differences between groups (Fig. 4C).

**Figure 4.**
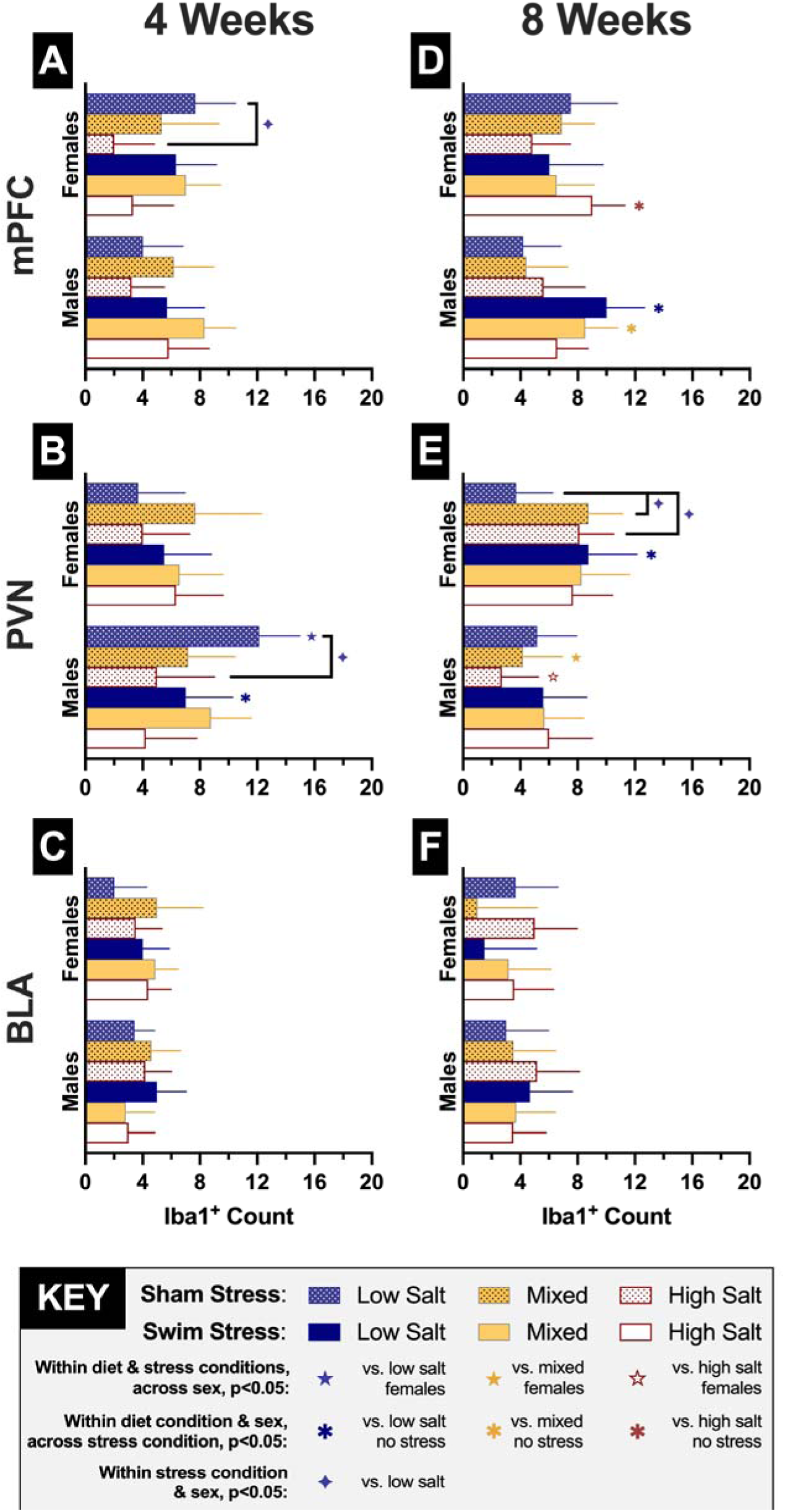
Counts of Iba1^+^ cells in three different brain regions. Iba1^+^ counts per hemisphere in 4 wk (panels A-C) and 8 wk (panels D-F) mice, within the medial prefrontal cortex (mPFC; panels A, D), paraventricular nucleus of the hypothalamus (PVN; panels B, E), and basolateral amygdala (BLA; panels C, F). Mice assigned to low salt diet (0.4% NaCl, w/w) are shown in blue bars. Mice assigned to high salt diet (4.0% NaCl) are displayed in white-filled red bars. Mixed cages given free choice access to both low and high salt diets are graphed in grey-bordered yellow bars. Dotted bars are for sham stress mice, clear bars are for swim stress mice. Ns=3-10, 3-8, 2-10 in mPFC, PVN, and BLA, respectively, within each sex/diet/stress condition; for exact numbers, see Supplemental Tables S12, S13, S14, respectively. Data are graphed as EM means ± 95% CI. In alphabetical panel order, top to bottom, ⍰indicate p=0.019, 0.016, 0.017, 0.044 vs. indicated group within same sex/stress/time period. 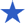 indicates p<0.001 vs. 4 wk sham stress low salt females. 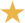 indicates p=0.015 vs. 8 wk sham stress mixed females. 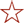 indicates p=0.003 vs. 8 wk sham stress high salt females. In alphabetical panel order, ⍰indicates p=0.022, 0.003, 0.021 vs. low salt sham stress of same sex and time period. ⍰indicates p=0.021 vs. high salt sham stress of same sex and time period. ⍰indicates p=0.031 vs. high salt sham stress of same sex and time period.

**Table 4.** Two-way RM GLMs of 4 wk Iba1^+^ counts in three brain regions.

| **4 Week Iba1<sup>+</sup> Counts** |  |  |  |
| --- | --- | --- | --- |
| **mPFC** | **F statistic** | **p** | **$\eta_p^2$** |
| Stress | $F_{(1,67)}=2.768$ | 0.101 | 0.040 |
| Sex | $F_{(1,67)}=0.103$ | 0.749 | 0.002 |
| **Diet** | **$F_{(2,67)}=5.350$** | **0.007** | **0.138** |
| Stress × Sex | $F_{(1,67)}=0.963$ | 0.330 | 0.014 |
| Stress × Diet | $F_{(2,67)}=0.521$ | 0.596 | 0.015 |
| Sex × Diet | $F_{(2,67)}=2.336$ | 0.105 | 0.065 |
| Stress × Sex × Diet | $F_{(2,67)}=0.216$ | 0.807 | 0.006 |
| **PVN** | **F statistic** | **p** | **$\eta_p^2$** |
| Stress | $F_{(1,59)}=0.046$ | 0.831 | 0.001 |
| Sex | $F_{(1,59)}=3.133$ | 0.082 | 0.050 |
| *Diet* | $F_{(2,59)}=2.599$ | *0.083* | *0.081* |
| Stress × Sex | $F_{(1,59)}=1.561$ | 0.216 | 0.026 |
| Stress × Diet | $F_{(2,59)}=0.585$ | 0.560 | 0.019 |
| *Sex × Diet* | $F_{(2,59)}=3.012$ | *0.057* | *0.093* |
| Stress × Sex × Diet | $F_{(2,59)}=2.067$ | 0.136 | 0.065 |
| **BLA** | **F statistic** | **p** | **$\eta_p^2$** |
| Stress | $F_{(1,59)}=0.155$ | 0.695 | 0.003 |
| Sex | $F_{(1,59)}=0.050$ | 0.824 | 0.001 |
| Diet | $F_{(2,59)}=0.482$ | 0.620 | 0.016 |
| Stress x Sex | $F_{(1,59)}=1.374$ | 0.246 | 0.023 |
| Stress x Diet | $F_{(2,59)}=1.922$ | 0.155 | 0.061 |
| Sex x Diet | $F_{(2,59)}=1.421$ | 0.250 | 0.046 |
| Stress x Sex x Diet | $F_{(2,59)}=0.204$ | 0.816 | 0.007 |

Analyses of 8 wk time period Iba1^+^ cell counts in the mPFC reported a significant three-way sex × stress × diet interaction (Table 5). Pairwise comparisons indicated that Iba1^+^ cell counts were increased in swim stress 8 wk HS females compared to sham stress 8 wk HS females (Fig. 4D). In contrast, swim stress 8 wk males in the LS and mixed diet conditions displayed elevated Iba1^+^ cell counts compared to sham stress males in the respective diet conditions (Fig. 4D).

**Table 5.** Two-way RM GLMs of 8 wk Iba1^+^ counts in three brain regions.

| 8 Week Iba1<sup>+</sup> Counts |  |  |  |
| --- | --- | --- | --- |
| mPFC | F statistic | p | $\eta_p^2$ |
| Stress | $F_{(1,62)}=7.690$ | 0.007 | 0.110 |
| Sex | $F_{(1,62)}=0.098$ | 0.756 | 0.002 |
| Diet | $F_{(2,62)}=0.095$ | 0.910 | 0.003 |
| Stress × Sex | $F_{(1,62)}=3.272$ | 0.075 | 0.050 |
| Stress × Diet | $F_{(2,62)}=0.076$ | 0.927 | 0.002 |
| Sex × Diet | $F_{(2,62)}=0.175$ | 0.840 | 0.006 |
| **Stress × Sex × Diet** | **$F_{(2,62)}=4.052$** | **0.022** | **0.116** |
| PVN | F statistic | p | $\eta_p^2$ |
| Stress | $F_{(1,60)}=3.573$ | 0.064 | 0.056 |
| **Sex** | **$F_{(1,60)}=10.51$** | **0.002** | **0.149** |
| Diet | $F_{(2,60)}=0.399$ | 0.673 | 0.013 |
| Stress × Sex | $F_{(1,60)}=0.054$ | 0.817 | 0.001 |
| Stress × Diet | $F_{(2,60)}=0.596$ | 0.554 | 0.019 |
| Sex × Diet | $F_{(2,60)}=1.168$ | 0.318 | 0.037 |
| Stress × Sex × Diet | $F_{(2,60)}=2.346$ | 0.104 | 0.073 |

| BLA | F statistic | p | $\eta_p^2$ |
| --- | --- | --- | --- |
| Stress | $F_{(1,61)}=0.052$ | 0.820 | 0.001 |
| Sex | $F_{(1,61)}=1.125$ | 0.293 | 0.018 |
| Diet | $F_{(2,61)}=1.069$ | 0.350 | 0.034 |
| Stress x Sex | $F_{(1,61)}=0.095$ | 0.759 | 0.002 |
| Stress x Diet | $F_{(2,61)}=0.820$ | 0.445 | 0.026 |
| Sex x Diet | $F_{(2,61)}=0.285$ | 0.753 | 0.009 |
| Stress x Sex x Diet | $F_{(2,61)}=0.875$ | 0.422 | 0.028 |

PVN Iba1^+^ cell counts did not have any three-nor two-way interactions, but a significant main effect of sex was found (Table 5). Within sham stress 8 wk females, both mixed and HS females had higher Iba1^+^ cell counts than LS females (Fig. 4E). For swim stress 8 wk females, while no diet differences were observed, LS females that experienced swim stress exhibited more Iba1^+^ cell counts than sham stress 8 wk females (Fig. 4E). For sham stress 8 wk mice, male mixed and HS Iba1^+^ cell counts were significantly lower than counts in sham stress 8 wk females eating the same respective diets. As with the BLA in 4 wk mice, both omnibus statistics and pairwise comparisons for 8 wk BLA Iba1^+^ cell counts were negative (Table 5, Fig. 4F).

## Discussion

To expand the translatability and scope of dietary salt research, we explored here how access to HS diet affected adult female and male consumption patterns, active and passive coping behaviors, and a marker of neuroinflammation in three separate stress-responsive brain regions. Our choice access diet condition revealed mice voluntarily ate more salty food over time, with females exhibiting this pattern sooner than males. Passive, but not active, stress coping behaviors were sex-specifically shifted in mixed mice, as were 4 wk Iba1^+^ counts in sham stress mice. Only in the 8 wk condition were sex- and diet-dependent augmenting effects of stress on Iba1^+^ counts revealed.

The modest sizes of the salt coping behavior ^16,55,62,93^ and salt Iba1 ^16–21,64^ literature groupings in rodents mean our findings contribute to multiple dimensions. Most studies have assessed only single sexes ^18,20,64,93^, including our own ^16,62^. Others did not state the sex(es) used, or were admirably sex inclusive yet didn’t analyze sex differences ^17,19,21^. Such studies, while informative, can conceal biobehavioral outcomes relevant to about half of the world’s population. Two sex inclusive studies began excess salt diets at weaning (i.e., 21 days old; ^17,55^), revealing influential impacts of eating salty food during adolescence, and raising questions regarding how similar findings might be if the biobehavioral complexities of adolescence were omitted. Here, we focused on manipulating dietary salt across sexes starting in adulthood, and incorporated sex as a factor for all our analyses.

With so few studies of rodents across sexes in these salt literature groupings, evaluations of cross sex body weights and consumption are likewise limited. Fan and colleagues ^17^ reported no effect of diet on either measure in mice, but did not analyze sex effects. However, Lago and coworkers found that despite food intake not differing across diet, male rats eating salty food weighed less than control males near the end of their 6 wk experiment, whereas no such diet effect occurred in females ^55^. Neither of the latter controlled for sexually dimorphic body weights when analyzing diet intake ^17,55^, nor did we when studying salty food and fear conditioning ^63^. Aligning with previous findings ^17,55,63^, body weights of mice in the present study predominantly did not differ across diets in males nor females. Thus, we subsequently rectified our previous oversight by normalizing cage consumption to cage body weight.

To augment our research’s translatability, we implemented a free access, home cage choice condition of two diet formulations with ten-fold different salt levels to assess mice’s food preferences. In agreement with prior work ^17,55,63^, heightened salt levels did not affect overall normalized total food intake rates in a consistent manner across sexes. Across time periods, mixed females immediately consumed similar amounts of both LS and HS diets, whereas males initially preferred LS significantly more than HS chow. By the end of the 4 wk (males only) and 8 wk (females and males) periods, mice significantly preferred HS over LS food. We interpret these patterns as voluntary shifts in salty food preference that are not confounded by broader food overindulgence nor avoidance. Further, this voluntary consumption of more salt than biologically necessary - when accessible - both supports our hypothesis and reflects what is observed in people.

We did not observe our hypothesized differences in active coping behavior between LS and HS mice in either sex nor time period. It is therefore not surprising the LS vs. HS diet preference shifts across sexes assigned to the mixed condition likewise did not display active coping differences. Similarly, this makes the passive coping sex × diet interaction all the more unanticipated, given it seems driven by increases in mixed males, but decreases in mixed females. Likely, the implementation of salt loading facilitated the robust behavior effects we’ve observed previously ^16,62^, though others have reported no impact of salt loading on coping during a tail suspension test ^93^. Lago and colleagues were the first to report on acute stress coping behaviors across sexes, and the only amidst this small literature group to use salty food rather than salt loading ^55^. Their omnibus analyses revealed sex did not interact with diet to affect behavior, but that salt diet affected active coping responses in the swim stress ^55^. Graphs of their data illustrate heightened active coping accompanying an increased latency in shifting from an active to passive coping strategy ^55^. Our findings partially align with theirs, insofar as our data similarly suggest less pronounced coping strategy shifts in females compared to males. However, longer HS access in males accelerated strategy switching, contrasting with their findings ^55^. These discrepancies could be due to species differences, age at diet manipulation onset, higher dietary salt percentage, and/or diet manipulation duration ^55^. Though stress responsive behaviors investigated here were not as anticipated, they are nevertheless informative. These data provide the first such assessments in female rodents receiving adult-onset HS diet access, plus generate behavior data on our inaugural salt diet formulation choice procedure. Such experiments substantially expand the translatability of dietary salt research to people. Additionally, swim stressors served dual purposes by both facilitating coping analyses and adding an environmental stress manipulation for subsequent evaluations of Iba1 labeling.

Considering 4 wk mixed mice willingly consumed similar (females) if not greater (males) levels of HS vs LS food, one might expect that their Iba1^+^ counts would reflect those of HS mice. That such was not observed for sham stress females’ mPFCs, and sham stress males’ PVNs, suggests that even modest consumption of LS food might mitigate HS intake effects. However, for both these instances, HS lowered Iba1^+^ counts, suggestive of less neuroinflammation and thus potentially increased brain health (but see ^91,92^). Additional studies are needed to explore this possibility. Relatedly, swim stress in 4 wk male LS mice unexpectedly reduced PVN Iba1^+^ counts, but such swim effects were absent in HS access males. Thus, HS access could disrupt acute stress-elicited anti-neuroinflammatory processes, a line of future inquiry that would provide broad NCD-relevant information.

In contrast to 4 wk Iba1^+^ data, counts of Iba1^+^ cells in 8 wk mice at least partially supported our hypotheses. HS access in 8 wk sham stress mice preferentially increased female PVN Iba1^+^ counts, both compared to LS females and to HS access males. Further, 8 wk LS females exhibited increased PVN Iba1^+^ counts due to stress, but HS access prevented this stress augmentation in 8 wk females. Such time-, sex-, and stress-specific effects indicate that excess salt’s impacts on neuroinflammation are far from straightforward, but instead likely vary based upon circulating levels of sex hormones in conjunction with stressor encounters and duration of HS access. Supporting this complexity were 8 wk mPFC Iba1^+^ counts that revealed stress-selective diet and sex effects. In 8 wk females, swim stress exposure only elevated mPFC Iba1^+^ counts in HS, a portion of what we hypothesized. Contrastingly, and unexpectedly, swim stress selectively enhanced 8 wk male mPFC Iba1^+^ counts in mice with LS access. Collectively, our Iba1 findings provide critical evidence that, under certain sex and stress conditions, access to salty food can attenuate a neuroinflammatory marker. This contrasts with all previously published findings, including our own, reporting that excess salt intake increases or does not affect Iba1 measures across 10 total brain regions ^16–21,64^.

Some of the incongruencies between the current findings and existing salt Iba1 literature may be because our salt manipulations did not commence until adulthood rather than at weaning ^17^, didn’t extend beyond 8 wks ^18,19^, did not use salt loading ^16,64^ nor supplement salty food with salt loading ^17–19,21^, and/or that we used 4% instead of 8% NaCl chow ^18,20,21^. We intentionally used only HS chow as our diet manipulation to improve translatability. First, people generally do not drink meaningful amounts of salt in beverages unless the goal is emesis. More frequently, criticisms arise regarding the salt percentage in chow. While 4% might initially seem like a large amount of salt in food, the ability of mice to excrete sodium 8 times better than humans must be factored in ^94^. Setting aside additional factors that make mice still more capable of tolerating high sodium levels versus humans ^94^, a simple calculation indicates that this diet could roughly approximate 0.5% NaCl (w/w) in human foods. This translates to 0.20% sodium (w/w).

Unprocessed foods range in their natural sodium levels (all w/w), from low amounts in long grain brown rice (0.03%), to moderate levels in celery (0.10%) and eggs (0.12%), topping out in sea-based products like shrimp (0.48%) ^95^. Some processed foods, like Oreo® (0.38%, w/w) ^96^ and white bread (0.49%, w/w) ^95^, have comparable levels. Other processed foods can be much higher, like yellow mustard (1.1%, w/w) ^95^ and Kraft® Singles American Slices (1.09%) ^97^.

Across several European countries, ham and bacon vary in salt amounts from 1.7-3.2%, w/w ^98^. As evidenced by this handful of examples, the ranges of sodium/salt levels that people ingest are widespread (see reviews ^49–51)^. Therefore, our use of HS chow in mice is actually a modest dietary salt manipulation in this context, maximizing insights into salt’s brain and behavior effects applicable to a majority of people.

Combined, our study has: 1) introduced a diet formulation choice procedure into rodent research; 2) discovered that across sexes mice voluntarily choose to eat saltier food than biologically necessary, akin to humans; 3) found HS consumption can decrease Iba1^+^ counts across sexes in the absence of stress; and, 4) demonstrated how this neuroinflammatory marker varies across brain regions depending upon combinations of biological and environmental factors. A limitation of this investigation was group housing of mice. This was done to minimize potential stress confounds resulting from singly housing ^99–101^ (but see ^102,103^), but resulted in the inability to directly link individual consumption of food/salt to individual Iba1^+^ counts, as consumption was quantified per cage. The advantage of this approach, though, was reducing the possibility that our Iba1^+^ counts were confounded by different housing conditions. For each cage assigned to a given diet manipulation, at least one mouse underwent swim stress, and at least one mouse underwent sham stress. In future studies, we will evaluate singly housed mice’s eating choices under a mixed salt diet condition. Iba1 labeling itself is limited in that it cannot distinguish between different forms of activated microglia ^91^, plus some health conditions can suppress Iba1 labeling while neuroinflammation persists (see thorough review by ^92^). This also provides an alternative interpretation for our HS-induced reductions in Iba1. Relatedly, another limitation is that, due to methodological constraints, we could only analyze hemispheres from a subset of mice that were utilized for consumption and behavior analyses, reducing the *a priori* planned power of our Iba1^+^ queries. We are curious to learn how much of our findings replicate in the hands of others performing similar investigations. Even in our own hands, our data did not always replicate. An example is normalized food intake between mixed male 4 wk and 8 wk mice. While by the end of the 4 wk period in the 4 wk experiment, mixed males significantly reversed their diet preference from LS to HS, it took the 8 wk mice until their 8^th^ wk to exhibit the same reversal. We suspect this is due to group housing of our mice, meaning individual preferences of mixed mice within the same cage could shift the apparent cage preference for diets, and also influence the rate at which each cage achieved reversal. Again, group housing still afforded benefits for our experimental design, but future experiments are needed to assess the degree of variation between individual mice regarding shifting salty food preferences. Future experiments evaluating the influences of different stressors during, rather than after, dietary salt manipulations would be informative regarding how salty food choices change during or after stress. Evaluations of stressors lasting longer than 6 min would also provide more insights regarding how HS access influences stress coping behaviors. The stress-enhanced adrenocorticotropic hormone both facilitates hormonal responses to high arousal states and stimulates release of the sodium retention-promoting hormone aldosterone ^104,105^. Assessments of these and other biological markers connecting stress and salt intake could uncover molecular links between stress responsivity and eating behaviors that contribute to, and might even serve as biomarkers for, NCDs. Consequently, the present findings open a plethora of investigative opportunities into the intersecting dimensions of biological sex, environmental stress, salty food preferences, and neuroinflammation.

## Acknowledgements

This project was supported by Kent State University and, in part, by a Distinguished Dissertation Award from Kent State University’s Healthy Communities Research Institute (HCRI) and a Judie Fall Lasser Graduate Psychology Research Award from Kent State University’s Department of Psychological Sciences, both to JNB. JNB was also supported by a Roy S. Lilly Graduate Student Scholarship from Kent State University’s Department of Psychological Sciences, and a Lillian Friedman Graduate Student Scholarship from Kent State University. Content is the authors’ responsibility and does not represent the views of HCRI. The authors express their thanks to their expert veterinarians and vivarium personnel, and gratefully acknowledge all the mice used in these experiments.

