## Supplemental Materials for "Voluntary eating of saltier food by mice and acute stress each abrogate reductions in a neuroinflammatory marker across sexes"

*Corresponding author

Lee M. Gilman, Ph.D.

600 Hilltop Dr.

144 Kent Hall

Kent State University

Kent, OH 44242 USA

+1 (330) 672-2201

Supplemental Table S1. Two-way repeated measures (RM) general linear models (GLMs) of individual mice’s body weights.

| **Body Weights** | **F statistic** | **p** | **Partial η^2^ (η_p_^2^)** |
| --- | --- | --- | --- |
| **4 Weeks** |  |  |  |
| **Week** | **F_(3.082,299.0)_=15.87** | **<0.001** | **0.141** |
| **Sex** | **F_(1,97)_=180.7** | **<0.001** | **0.651** |
| Diet | F_(2,97)_=1.564 | 0.214 | 0.031 |
| Week × Sex | F_(3.082,299.0)_=1.499 | 0.214 | 0.015 |
| Week × Diet | F_(6.165,299.0)_=1.223 | 0.294 | 0.025 |
| Sex × Diet | F_(2,97)_=1.270 | 0.285 | 0.026 |
| Week × Sex × Diet | F_(6.165,299.0)_=0.349 | 0.914 | 0.007 |
| **8 Weeks** |  |  |  |
| Week | F_(3.276,285.0)_=50.17 | <0.001 | 0.366 |
| Sex | F_(1,87)_=226.3 | <0.001 | 0.722 |
| Diet | F_(2,87)_=2.502 | 0.088 | 0.054 |
| **Week × Sex** | **F_(3.276,285.0)_=7.317** | **<0.001** | **0.078** |
| Week × Diet | F_(6.552,285.0)_=1.397 | 0.211 | 0.031 |
| Sex × Diet | F_(2,87)_=0.247 | 0.782 | 0.006 |
| Week × Sex × Diet | F_(6.552,285.0)_=1.760 | 0.101 | 0.039 |

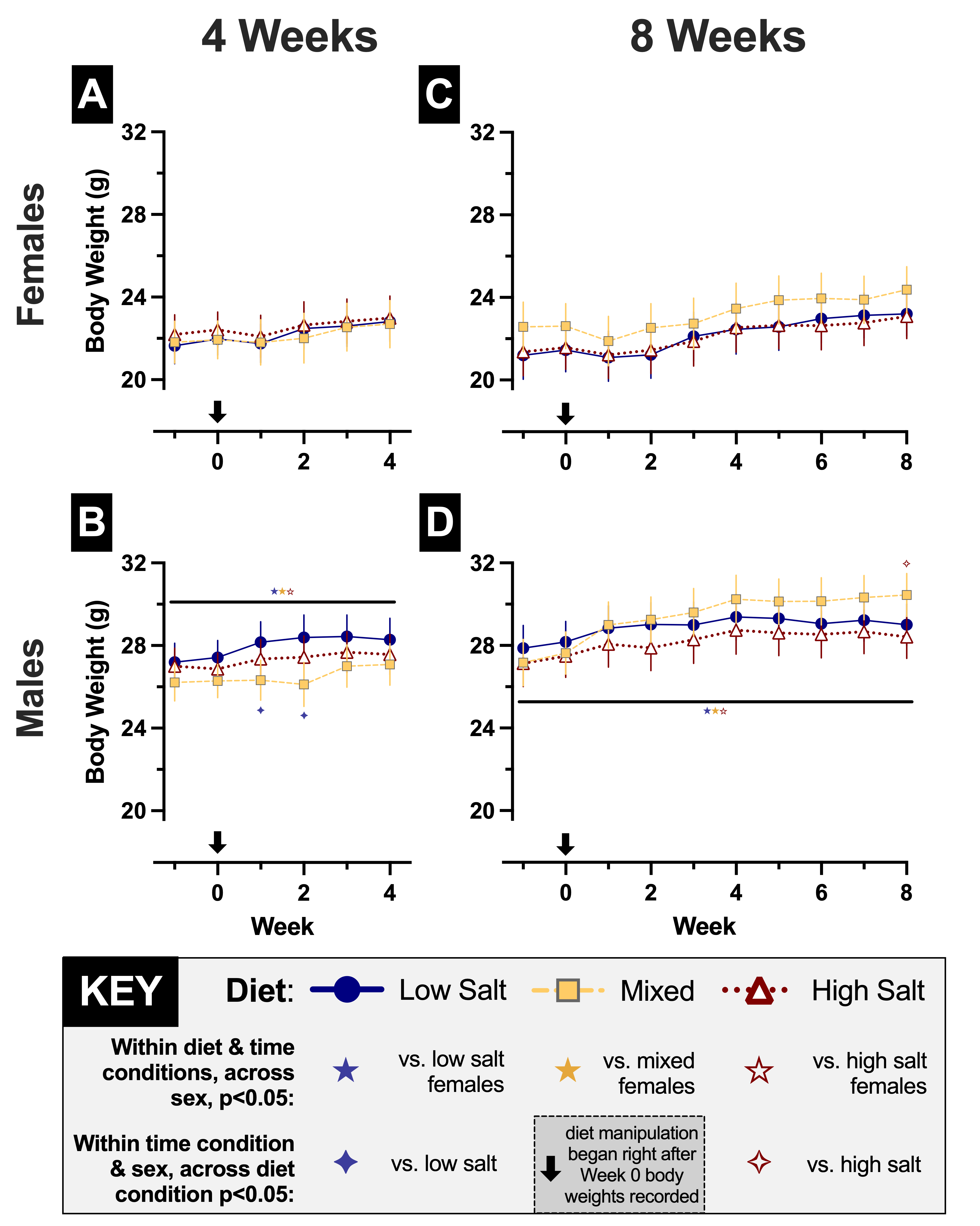

Supplemental Figure S1. Body weights of individual mice within same sex cages assigned to a diet condition. Body weights were recorded weekly for 4 wk (panels A-B) and 8 wk (panels C-D) mice, starting at week −1. Female weights are shown in panels A,C; male weights are in panels B, D. Mice assigned to low salt diet (0.4% NaCl, w/w) are shown in blue circles with solid lines. Mice assigned to high salt diet (4.0% NaCl) are displayed in white-filled red triangles with dotted lines. Mixed mice with free choice access to both low and high salt diets are graphed in grey-bordered yellow squares with dashed lines. Ns=14-19 within each sex/diet/time period; for exact numbers, see Supplemental Table S2. Data are graphed as estimated marginal means (EM means) ± 95% confidence interval (CI). Left to right, ✦indicates p=0.029, 0.012 vs. same sex low salt group at same week. ✧indicates p=0.023 vs. same sex high salt group at same week. Five point stars indicate p<0.001 compared with female ★low salt, ★mixed, and ☆high salt mice in same week and time period; significance applies to all weeks covered by horizontal black line. Down arrow indicates diet manipulation onset, commencing immediately after Week 0 body weights were recorded.

Supplemental Table S2. Numbers of individual mice graphed in Fig. S1.

| **Time Period** | **Sex** | **Diet Manipulation** | | |
| --- | --- | --- | --- | --- |
|  |  | **Low Salt** | **Mixed** | **High Salt** |
| **4 wks** | **Female** | 19 | 14 | 16 |
|  | **Male** | 17 | 18 | 19 |
| **8 wks** | **Female** | 16 | 16 | 16 |
|  | **Male** | 17 | 16 | 16 |

Supplemental Table S3. Two-way RM GLMs of baseline (Week 0) normalized food/day and normalized water/day.

| **Normalized Week 0** | **F statistic** | **p** | **η_p_^2^** |
| --- | --- | --- | --- |
| **Food/Day** |  |  |  |
| Time | F_(1,65)_=0.037 | 0.848 | 0.001 |
| **Sex** | **F_(1,65)_=5.807** | **0.019** | **0.082** |
| Diet | F_(2,65)_=0.187 | 0.830 | 0.006 |
| Time × Sex | F_(1,65)_=0.040 | 0.842 | 0.001 |
| Time × Diet | F_(2,65)_=0.670 | 0.515 | 0.020 |
| Sex × Diet | F_(2,65)_=0.030 | 0.971 | 0.001 |
| Time × Sex × Diet | F_(2,65)_=0.152 | 0.859 | 0.005 |
| **Water/Day** |  |  |  |
| Time | F_(1,66)_=0.002 | 0.968 | 0.000 |
| **Sex** | **F_(1,66)_=17.85** | **<0.001** | **0.213** |
| Diet | F_(2,66)_=0.127 | 0.881 | 0.004 |
| Time × Sex | F_(1,66)_=0.237 | 0.628 | 0.004 |
| Time × Diet | F_(2,66)_=0.470 | 0.627 | 0.014 |
| Sex × Diet | F_(2,66)_=0.648 | 0.527 | 0.019 |
| Time × Sex × Diet | F_(2,66)_=0.215 | 0.807 | 0.006 |

Supplemental Table S4. Numbers of cages graphed in Figs. 1, S2, and 2.

| **Time Period** | **Sex** | **Diet Manipulation** | | |
| --- | --- | --- | --- | --- |
|  |  | **Low Salt** | **Mixed** | **High Salt** |
| **4 wks** | **Female** | 7 | 5 | 6 |
|  | **Male** | 7 | 7 | 7 |
| **8 wks** | **Female** | 6 | 6 | 7 |
|  | **Male** | 7 | 7 | 6 |

Supplemental Table S5. Pairwise comparison results for 4 wk data in Fig. 1.

| **Wk** | ***Females*** | **Mixed Total** | **High Salt** | **Mixed-LS** | **Mixed-HS** |
| --- | --- | --- | --- | --- | --- |
| 1 | **Low Salt** | 1.000 | 0.940 | **0.014** | **<0.001** |
|  | **Mixed Total** |  | 1.000 | **0.002** | **<0.001** |
|  | **High Salt** |  |  | **<0.001** | **<0.001** |
|  | **Mixed-LS** |  |  |  | 0.263 |
| 2 | **Low Salt** | 0.805 | 0.097 | 0.055 | **0.019** |
|  | **Mixed Total** |  | 1.000 | **<0.001** | **<0.001** |
|  | **High Salt** |  |  | **<0.001** | **<0.001** |
|  | **Mixed-LS** |  |  |  | 1.000 |
| 3 | **Low Salt** | 1.000 | **0.039** | **<0.001** | **0.001** |
|  | **Mixed Total** |  | 1.000 | **<0.001** | **<0.001** |
|  | **High Salt** |  |  | **<0.001** | **<0.001** |
|  | **Mixed-LS** |  |  |  | 1.000 |
| 4 | **Low Salt** | 0.166 | **0.004** | **<0.001** | **0.004** |
|  | **Mixed Total** |  | 1.000 | **<0.001** | **<0.001** |
|  | **High Salt** |  |  | **<0.001** | **<0.001** |
|  | **Mixed-LS** |  |  |  | 0.370 |
| **Wk** | ***Males*** | **Mixed Total** | **High Salt** | **Mixed-LS** | **Mixed-HS** |
| 1 | **Low Salt** | 1.000 | 1.000 | **0.028** | **<0.001** |
|  | **Mixed Total** |  | 1.000 | **0.001** | **<0.001** |
|  | **High Salt** |  |  | **0.029** | **<0.001** |
|  | **Mixed-LS** |  |  |  | **0.001** |
| 2 | **Low Salt** | 1.000 | 1.000 | 0.067 | **<0.001** |
|  | **Mixed Total** |  | 1.000 | **<0.001** | **<0.001** |
|  | **High Salt** |  |  | **0.006** | **<0.001** |
|  | **Mixed-LS** |  |  |  | 1.000 |
| 3 | **Low Salt** | 0.590 | 1.000 | **<0.001** | **0.032** |
|  | **Mixed Total** |  | 1.000 | **<0.001** | **<0.001** |
|  | **High Salt** |  |  | **<0.001** | **0.011** |
|  | **Mixed-LS** |  |  |  | 0.380 |
| 4 | **Low Salt** | 0.397 | 1.000 | **<0.001** | **0.002** |
|  | **Mixed Total** |  | 1.000 | **<0.001** | **<0.001** |
|  | **High Salt** |  |  | **<0.001** | **<0.001** |
|  | **Mixed-LS** |  |  |  | **0.004** |

Supplemental Table S6. Pairwise comparison results for 8 wk data in Fig. 1.

| **Wk** | ***Females*** | **Mixed Total** | **High Salt** | **Mixed-LS** | **Mixed-HS** |
| --- | --- | --- | --- | --- | --- |
| 1 | **Low Salt** | 1.000 | 1.000 | 0.147 | **0.002** |
|  | **Mixed Total** |  | 1.000 | **0.002** | **<0.001** |
|  | **High Salt** |  |  | **0.001** | **<0.001** |
|  | **Mixed-LS** |  |  |  | 1.000 |
| 2 | **Low Salt** | 1.000 | 0.134 | **0.024** | 0.085 |
|  | **Mixed Total** |  | 1.000 | **<0.001** | **0.003** |
|  | **High Salt** |  |  | **<0.001** | **<0.001** |
|  | **Mixed-LS** |  |  |  | 1.000 |
| 3 | **Low Salt** | 1.000 | 0.715 | **0.003** | 0.263 |
|  | **Mixed Total** |  | 1.000 | **<0.001** | **0.015** |
|  | **High Salt** |  |  | **<0.001** | **0.001** |
|  | **Mixed-LS** |  |  |  | 1.000 |
| 4 | **Low Salt** | 0.334 | 0.252 | **0.002** | **0.028** |
|  | **Mixed Total** |  | 1.000 | **<0.001** | **<0.001** |
|  | **High Salt** |  |  | **<0.001** | **<0.001** |
|  | **Mixed-LS** |  |  |  | 1.000 |
| 5 | **Low Salt** | 0.942 | 0.277 | **0.001** | 1.000 |
|  | **Mixed Total** |  | 1.000 | **<0.001** | **0.024** |
|  | **High Salt** |  |  | **<0.001** | **0.004** |
|  | **Mixed-LS** |  |  |  | 0.097 |
| 6 | **Low Salt** | 0.910 | 0.183 | **<0.001** | 0.405 |
|  | **Mixed Total** |  | 1.000 | **<0.001** | **0.004** |
|  | **High Salt** |  |  | **<0.001** | **<0.001** |
|  | **Mixed-LS** |  |  |  | **0.003** |
| 7 | **Low Salt** | 1.000 | 0.179 | **<0.001** | 0.354 |
|  | **Mixed Total** |  | 1.000 | **<0.001** | **0.009** |
|  | **High Salt** |  |  | **<0.001** | **<0.001** |
|  | **Mixed-LS** |  |  |  | **<0.001** |
| 8 | **Low Salt** | 0.912 | 0.504 | **<0.001** | 0.061 |
|  | **Mixed Total** |  | 1.000 | **<0.001** | **<0.001** |
|  | **High Salt** |  |  | **<0.001** | **<0.001** |
|  | **Mixed-LS** |  |  |  | **0.011** |
| **Wk** | ***Males*** | **Mixed Total** | **High Salt** | **Mixed-LS** | **Mixed-HS** |
| 1 | **Low Salt** | 1.000 | 0.732 | 0.599 | **<0.001** |
|  | **Mixed Total** |  | 1.000 | 0.150 | **<0.001** |
|  | **High Salt** |  |  | **0.007** | **<0.001** |
|  | **Mixed-LS** |  |  |  | **<0.001** |
| 2 | **Low Salt** | 1.000 | 0.769 | 0.243 | **0.003** |
|  | **Mixed Total** |  | 1.000 | **0.028** | **<0.001** |
|  | **High Salt** |  |  | **0.002** | **<0.001** |
|  | **Mixed-LS** |  |  |  | 1.000 |
| 3 | **Low Salt** | 1.000 | 1.000 | **0.008** | **0.005** |
|  | **Mixed Total** |  | 1.000 | **0.008** | **0.005** |
|  | **High Salt** |  |  | **<0.001** | **<0.001** |
|  | **Mixed-LS** |  |  |  | 1.000 |
| 4 | **Low Salt** | 1.000 | 0.112 | **<0.001** | **<0.001** |
|  | **Mixed Total** |  | 0.237 | **<0.001** | **<0.001** |
|  | **High Salt** |  |  | **<0.001** | **<0.001** |
|  | **Mixed-LS** |  |  |  | 1.000 |
| 5 | **Low Salt** | 1.000 | 0.693 | **<0.001** | **0.008** |
|  | **Mixed Total** |  | 1.000 | **<0.001** | **0.006** |
|  | **High Salt** |  |  | **<0.001** | **<0.001** |
|  | **Mixed-LS** |  |  |  | 1.000 |
| 6 | **Low Salt** | 1.000 | **0.033** | **<0.001** | **0.003** |
|  | **Mixed Total** |  | 0.108 | **<0.001** | **0.001** |
|  | **High Salt** |  |  | **<0.001** | **<0.001** |
|  | **Mixed-LS** |  |  |  | 0.241 |
| 7 | **Low Salt** | 1.000 | 0.220 | **<0.001** | **<0.001** |
|  | **Mixed Total** |  | 0.105 | **<0.001** | **<0.001** |
|  | **High Salt** |  |  | **<0.001** | **<0.001** |
|  | **Mixed-LS** |  |  |  | 0.222 |
| 8 | **Low Salt** | 1.000 | 0.357 | **<0.001** | **0.002** |
|  | **Mixed Total** |  | 0.673 | **<0.001** | **0.001** |
|  | **High Salt** |  |  | **<0.001** | **<0.001** |
|  | **Mixed-LS** |  |  |  | **0.005** |

Supplemental Table S7. Two-way RM GLMs of normalized water/day.

| **Normalized Water/Day** |  |  |  |
| --- | --- | --- | --- |
| **4 Weeks** | **F statistic** | **p** | **η_p_^2^** |
| **Week** | **F_(2.459,81.16)_=13.47** | **<0.001** | **0.290** |
| Sex | F_(1,33)_=2.001 | 0.167 | 0.057 |
| Diet | F_(2,33)_=22.88 | <0.001 | 0.581 |
| Week × Sex | F_(2.459,81.16)_=1.814 | 0.161 | 0.052 |
| Week × Diet | F_(4.919,81.16)_=1.639 | 0.160 | 0.090 |
| **Sex × Diet** | **F_(2,33)_=4.418** | **0.020** | **0.211** |
| Week × Sex × Diet | F_(4.919,81.16)_=1.911 | 0.103 | 0.104 |
| **8 Weeks** | **F statistic** | **p** | **η_p_^2^** |
| Week | F_(3.265,94.70)_=21.21 | <0.001 | 0.422 |
| Sex | F_(1,29)_=1.178 | 0.287 | 0.039 |
| **Diet** | **F_(2,29)_=19.27** | **<0.001** | **0.571** |
| **Week × Sex** | **F_(3.265,94.70)_=7.034** | **<0.001** | **0.195** |
| Week × Diet | F_(6.531,94.70)_=0.674 | 0.683 | 0.044 |
| Sex × Diet | F_(2,29)_=1.223 | 0.309 | 0.078 |
| Week × Sex × Diet | F_(6.531,94.70)_=1.174 | 0.326 | 0.075 |

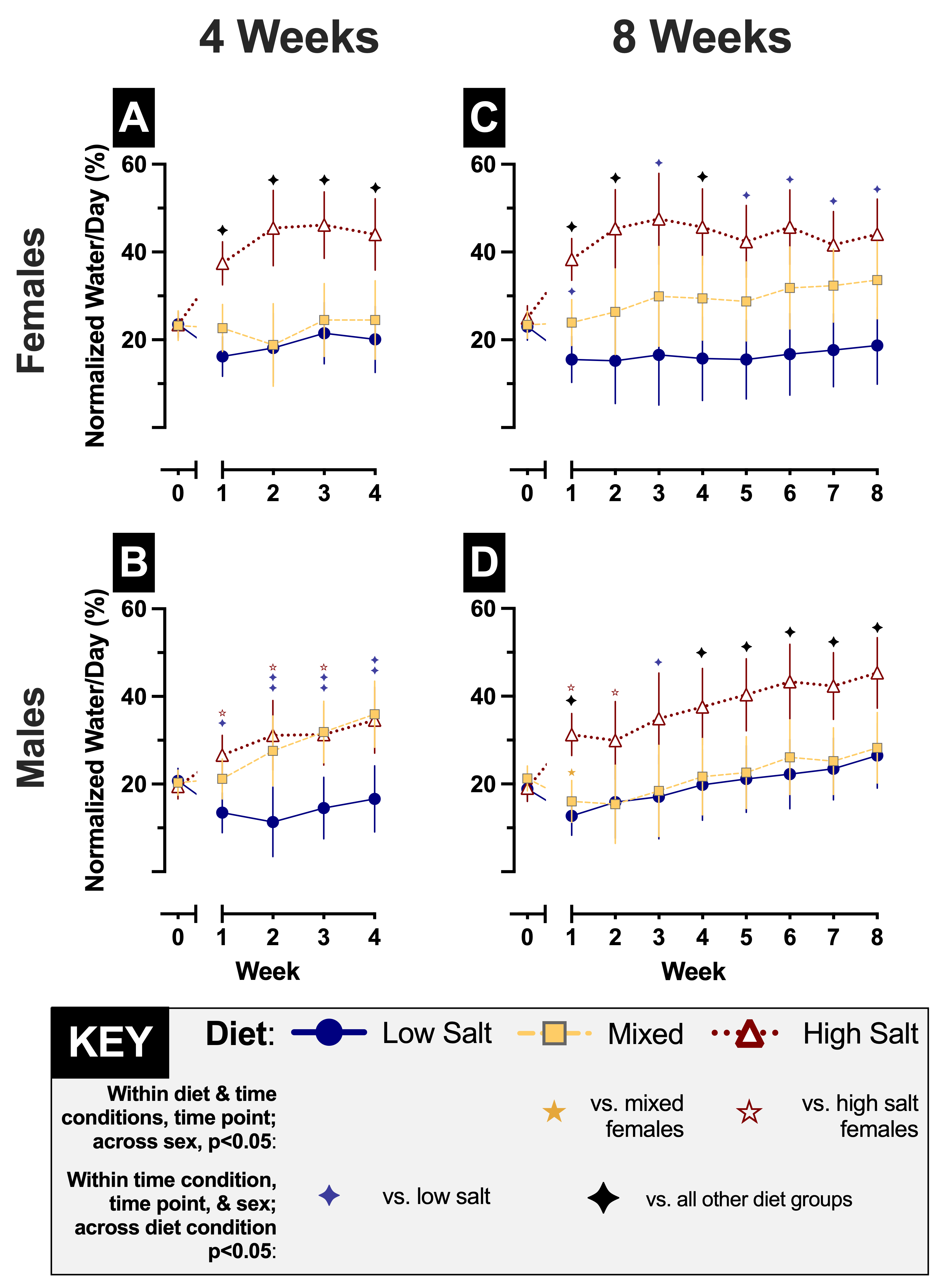

Supplemental Figure S2. Weekly rates of water consumed per day normalized to cage body weight. Normalized water/day intake was calculated weekly for 4 wk (panels A-B) and 8 wk (panels C-D) mice. Female cages are shown in panels A,C; male cages are in panels B, D. Cages assigned to low salt diet (0.4% NaCl, w/w) are shown in blue circles with solid lines. Cages assigned to high salt diet (4.0% NaCl) are displayed in white-filled red triangles with dotted lines. Mixed cages given free choice access to both low and high salt diets are graphed in grey-bordered yellow squares with dashed lines. Cage Ns=5-7 within each sex/diet/time period; for exact numbers, see Supplemental Table S4. Data are graphed as EM means ± 95% CI. Blue, yellow, and red diamonds indicate p<0.05 (see Tables S8 and S9 for 4 wk and 8 wk, respectively, exact p values) compared with ✦low salt, ✦mixed total, and ✧high salt cage consumption in same sex at same week within same time period. Black diamonds (✦) indicate p<0.05 versus all other points within same sex, week, and time period. Left to right, ☆indicates p=0.002, 0.018, 0.006, 0.041, 0.018 vs. high salt females in same week and time period. ★indicates p=0.031 vs. mixed females in same week and time period.

Supplemental Table S8. Pairwise comparison results for 4 wk data in Fig. S2.

| **Wk** | ***Females*** | **Mixed** | **High Salt** |
| --- | --- | --- | --- |
| 1 | **Low Salt** | 0.211 | **<0.001** |
|  | **Mixed** |  | **<0.001** |
| 2 | **Low Salt** | 1.000 | **<0.001** |
|  | **Mixed** |  | **<0.001** |
| 3 | **Low Salt** | 1.000 | **<0.001** |
|  | **Mixed** |  | **0.001** |
| 4 | **Low Salt** | 1.000 | **<0.001** |
|  | **Mixed** |  | **0.007** |
| **Wk** | ***Males*** | **Mixed** | **High Salt** |
| 1 | **Low Salt** | 0.060 | **<0.001** |
|  | **Mixed** |  | 0.285 |
| 2 | **Low Salt** | **0.018** | **0.003** |
|  | **Mixed** |  | 1.000 |
| 3 | **Low Salt** | **0.003** | **0.005** |
|  | **Mixed** |  | 1.000 |
| 4 | **Low Salt** | **0.002** | **0.005** |
|  | **Mixed** |  | 1.000 |

Supplemental Table S9. Pairwise comparison results for 8 wk data in Fig. S2.

| **Wk** | ***Females*** | **Mixed** | **High Salt** |
| --- | --- | --- | --- |
| 1 | **Low Salt** | 0.084 | **<0.001** |
|  | **Mixed** |  | **<0.001** |
| 2 | **Low Salt** | 0.326 | **<0.001** |
|  | **Mixed** |  | **0.019** |
| 3 | **Low Salt** | 0.300 | **<0.001** |
|  | **Mixed** |  | 0.079 |
| 4 | **Low Salt** | 0.140 | **<0.001** |
|  | **Mixed** |  | **0.047** |
| 5 | **Low Salt** | 0.125 | **<0.001** |
|  | **Mixed** |  | 0.088 |
| 6 | **Low Salt** | 0.078 | **<0.001** |
|  | **Mixed** |  | 0.098 |
| 7 | **Low Salt** | **0.049** | **<0.001** |
|  | **Mixed** |  | 0.311 |
| 8 | **Low Salt** | 0.062 | **<0.001** |
|  | **Mixed** |  | 0.250 |
| **Wk** | ***Males*** | **Mixed** | **High Salt** |
| 1 | **Low Salt** | 0.924 | **<0.001** |
|  | **Mixed** |  | **<0.001** |
| 2 | **Low Salt** | 1.000 | 0.074 |
|  | **Mixed** |  | 0.076 |
| 3 | **Low Salt** | 1.000 | **0.046** |
|  | **Mixed** |  | 0.087 |
| 4 | **Low Salt** | 1.000 | **0.014** |
|  | **Mixed** |  | **0.039** |
| 5 | **Low Salt** | 1.000 | **0.004** |
|  | **Mixed** |  | **0.012** |
| 6 | **Low Salt** | 1.000 | **0.003** |
|  | **Mixed** |  | **0.020** |
| 7 | **Low Salt** | 1.000 | **0.002** |
|  | **Mixed** |  | **0.009** |
| 8 | **Low Salt** | 1.000 | **0.004** |
|  | **Mixed** |  | **0.014** |

Supplemental Table S10. Pairwise comparison results for 8 wk data in Fig. 2.

| **Wk** | ***Females*** | **Mixed** | **High Salt** |
| --- | --- | --- | --- |
| 1 | **Low Salt** | **0.001** | **<0.001** |
|  | **Mixed** |  | **<0.001** |
| 2 | **Low Salt** | **<0.001** | **<0.001** |
|  | **Mixed** |  | **<0.001** |
| 3 | **Low Salt** | **<0.001** | **<0.001** |
|  | **Mixed** |  | **<0.001** |
| 4 | **Low Salt** | **<0.001** | **<0.001** |
|  | **Mixed** |  | **<0.001** |
| 5 | **Low Salt** | **<0.001** | **<0.001** |
|  | **Mixed** |  | **0.002** |
| 6 | **Low Salt** | **<0.001** | **<0.001** |
|  | **Mixed** |  | **<0.001** |
| 7 | **Low Salt** | **<0.001** | **<0.001** |
|  | **Mixed** |  | **<0.001** |
| 8 | **Low Salt** | **<0.001** | **<0.001** |
|  | **Mixed** |  | **<0.001** |
| **Wk** | ***Males*** | Mixed | High Salt |
| 1 | **Low Salt** | 0.051 | **<0.001** |
|  | **Mixed** |  | **<0.001** |
| 2 | **Low Salt** | **0.010** | **<0.001** |
|  | **Mixed** |  | **<0.001** |
| 3 | **Low Salt** | **0.004** | **<0.001** |
|  | **Mixed** |  | **<0.001** |
| 4 | **Low Salt** | **<0.001** | **<0.001** |
|  | **Mixed** |  | **<0.001** |
| 5 | **Low Salt** | **<0.001** | **<0.001** |
|  | **Mixed** |  | **<0.001** |
| 6 | **Low Salt** | **<0.001** | **<0.001** |
|  | **Mixed** |  | **<0.001** |
| 7 | **Low Salt** | **<0.001** | **<0.001** |
|  | **Mixed** |  | **<0.001** |
| 8 | **Low Salt** | **<0.001** | **<0.001** |
|  | **Mixed** |  | **<0.001** |

Supplemental Table S11. Numbers of individual mice graphed in Fig. 3.

| **Time Period** | **Sex** | **Diet Manipulation** | | |
| --- | --- | --- | --- | --- |
|  |  | **Low Salt** | **Mixed** | **High Salt** |
| **4 wks** | **Female** | 8 | 8 | 7 |
|  | **Male** | 9 | 9 | 10 |
| **8 wks** | **Female** | 7 | 8 | 8 |
|  | **Male** | 9 | 8 | 7 |

Supplemental Table S12. Medial prefrontal cortex hemispheres immunolabeled for Iba1 graphed in Fig. 4.

|  | **Time Period** | **Sex** | **Diet Manipulation** | | |
| --- | --- | --- | --- | --- | --- |
|  |  |  | **Low Salt** | **Mixed** | **High Salt** |
| **Sham Stress** | **4 wks** | **Female** | 6 | 3 | 6 |
|  |  | **Male** | 6 | 6 | 9 |
|  | **8 wks** | **Female** | 4 | 8 | 6 |
|  |  | **Male** | 6 | 5 | 5 |
| **Swim Stress** | **4 wks** | **Female** | 6 | 8 | 6 |
|  |  | **Male** | 7 | 10 | 6 |
|  | **8 wks** | **Female** | 3 | 6 | 8 |
|  |  | **Male** | 6 | 8 | 9 |

Supplemental Table S13. Paraventricular nucleus of the hypothalamus hemispheres immunolabeled for Iba1 graphed in Fig. 4.

|  | **Time Period** | **Sex** | **Diet Manipulation** | | |
| --- | --- | --- | --- | --- | --- |
|  |  |  | **Low Salt** | **Mixed** | **High Salt** |
| **Sham Stress** | **4 wks** | **Female** | 6 | 3 | 6 |
|  |  | **Male** | 8 | 6 | 4 |
|  | **8 wks** | **Female** | 7 | 8 | 8 |
|  |  | **Male** | 6 | 6 | 7 |
| **Swim Stress** | **4 wks** | **Female** | 6 | 7 | 6 |
|  |  | **Male** | 6 | 8 | 5 |
|  | **8 wks** | **Female** | 4 | 4 | 6 |
|  |  | **Male** | 5 | 6 | 5 |

Supplemental Table S14. Basolateral amygdala hemispheres immunolabeled for Iba1 graphed in Fig. 4.

|  | **Time Period** | **Sex** | **Diet Manipulation** | | |
| --- | --- | --- | --- | --- | --- |
|  |  |  | **Low Salt** | **Mixed** | **High Salt** |
| **Sham Stress** | **4 wks** | **Female** | 4 | 2 | 6 |
|  |  | **Male** | 10 | 5 | 6 |
|  | **8 wks** | **Female** | 6 | 3 | 6 |
|  |  | **Male** | 6 | 6 | 6 |
| **Swim Stress** | **4 wks** | **Female** | 6 | 8 | 8 |
|  |  | **Male** | 5 | 5 | 6 |
|  | **8 wks** | **Female** | 4 | 6 | 7 |
|  |  | **Male** | 6 | 7 | 10 |
